# Localization of four class I glutaredoxins in the cytosol and the secretory pathway and characterization of their biochemical diversification

**DOI:** 10.1101/2023.09.01.555924

**Authors:** Michelle Schlößer, Anna Moseler, Yana Bodnar, Maria Homagk, Stephan Wagner, Luca Pedroletti, Manuela Gellert, José M. Ugalde, Christopher H. Lillig, Andreas J. Meyer

## Abstract

Class I glutaredoxins (GRXs) are catalytically active oxidoreductases and considered key proteins mediating reversible glutathionylation and deglutathionylation of protein thiols during development and stress responses. To narrow in on putative target proteins, it is mandatory to know the subcellular localization of the respective GRXs and to understand their catalytic activities and putative redundancy between isoforms in the same compartment. We show that GRXC1 and GRXC2 are cytosolic proteins with GRXC1 being attached to membranes through myristoylation. GRXC3 and GRXC4 are identified as type II membrane proteins along the early secretory pathway with their enzymatic function on the luminal side. Comparison of all four studied GRXs for their oxidoreductase function highlights biochemical diversification with GRXC1 and GRXC2 being better reductants than GRXC3 and GRXC4 with bis(2-hydroxyethyl) disulfide and oxidized roGFP2 as substrates. *Vice versa*, GRXC3 and GRXC4 are better oxidants of reduced roGFP2 in the reverse reaction. Analysis of electrostatic surface potentials mirrors the phylogenetic classification of class I GRXs but cannot fully account for the observed kinetic differences in their interaction with roGPF2. Despite localization of two class I GRXs each in the cytosol and the endomembrane system, the respective double null mutants are viable without obvious phenotypes.

**Summary statement:** We identify Arabidopsis glutaredoxins GRXC3 and GRXC4 as type II membrane proteins in the secretory pathway and GRXC1 as attached to membranes through N-terminal myristoylation. Cytosolic GRXC1 and GRXC2 and luminal GRXC3 and GRXC4 display distinct biochemical properties in their redox activities.

## 1 | INTRODUCTION

In plants, stress-related responses and several developmental processes are accompanied with formation of reactive oxygen species (ROS) and subsequent translation of this primary oxidation to changes in protein thiol oxidation as a key step in stress signaling pathways (A. J. Meyer et al., 2021; Mittler et al., 2022). Detoxification of H_2_O_2_ through the ascorbate-glutathione cycle causes a transitory oxidation of glutathione with the oxidation of reduced glutathione (GSH) to glutathione disulfide (GSSG) (Foyer et al., 1994). Oxidative stress thus leads to a decrease of the glutathione redox potential (*E*_GSH_), which may be translated to changes in the redox status of protein thiols downstream (Deponte & Lillig, 2015; A. J. Meyer et al., 2007). The respective redox equilibration between glutathione and protein thiols may be mediated by glutaredoxins (GRXs) as thiol switch operators (Gutscher et al., 2008; A. J. Meyer, 2008; Ströher & Millar, 2012).

Glutaredoxins are ubiquitous proteins that belong to the thioredoxin (TRX) superfamily, members of which share the TRX fold as their defining three-dimensional structure. The TRX fold consists of a central four-stranded β-sheet with three flanking α-helices and the active site motif (Berndt et al., 2015; Pan & Bardwell, 2006). In *Arabidopsis thaliana*, GRXs form a large family with 33 members grouped in four distinct classes (I-IV) (Couturier et al., 2009; Y. Meyer et al., 2009). Members of the more distant class IV are multidomain proteins with an N-terminal GRX domain(Couturier et al., 2009). Class III GRXs (syn. ROXYs) have been identified as interacting proteins of TGA transcription factors, which together with their first appearance in land plants and pronounced diversification along the evolution of higher plants led to speculation about redox-sensitive transcriptional regulation in plant development (Gutsche et al., 2017; Xing et al., 2005). So far, however, no oxidoreductase activity has been reported (Mrozek et al., 2023). Class II GRXs are characterized by a CGFS active site and a characteristic five-amino acid loop adjacent to the active site that is missing in other GRXs (Trnka et al., 2020). The loop causes an altered orientation of the thiol group of the cofactor glutathione such that a strong preference of class II GRXs for coordination of [2Fe–2S] clusters is generated. The ability to coordinate a [2Fe–2S] cluster and to deliver it to the respective apoproteins defines the functional role of these GRXs as Fe–S cluster transferases (Pedroletti et al., 2023; Trnka et al., 2020).

Despite the nomenclature of the entire protein family as GRXs, the only representatives with unequivocal oxidoreductase activity are class I GRXs (Begas et al., 2017; Liedgens et al., 2020; Mrozek et al., 2023; Trnka et al., 2020). Class I GRXs are evolutionary old proteins and widespread in archaea, bacteria and eukaryotes, but expanded during plants conquered the land (Alves et al., 2009; Couturier et al., 2009). In Arabidopsis, this class comprises five members named GRXC1-5 plus GRXS12, which is closely related to GRXC5 (Rouhier et al., 2004). Of these proteins, GRXC1 is present exclusively in eudicots, suggesting that it may have evolved along with the development of more complex signaling mechanism. A common feature of all six class I GRXs is their C(P/G/S)Y(C/S) active site motif that enables the proteins to catalyze reversible glutathionylation and deglutathionylation of protein thiols depending on the local *E*_GSH_ (Liedgens et al., 2020; Trnka et al., 2020). Although no unequivocal evidence for such a catalytic function has been shown for endogenous target proteins *in planta*, class I GRXs have been exploited as thiol switch operators for the engineered redox-sensitive GFP2 (roGFP2) to enable sensitive, dynamic readouts of *E*_GSH_ in different compartments of live cells (Gutscher et al., 2008; A. J. Meyer et al., 2007; Schwarzländer et al., 2008). With the exception of GRXS12, all class I GRXs contain two cysteine residues in their active sites interspaced by two other amino acids. Although the second cysteine is not involved in glutathionylation or deglutathionylation reactions, intramolecular disulfide within the active site can be formed (Couturier et al., 2013). It has been shown that the presence of the second cysteine influences the reactivity of the catalytic cysteine in poplar (*Populus trichocarpa*) *Pt*GRXC1 and *Pt*GRXC2, but not in *Pt*GRXC3 and *Pt*GRXC4, which may contribute to biochemical diversification. For *Pt*GRXC1 and *Pt*GRXC2, it has been reported that they are less efficient reductive catalysts compared to GRXC3 and GRXC4 (Couturier et al., 2013).

Glutathione is well established to play key roles in the response to a multitude of abiotic and biotic stresses (Noctor et al., 2012; Noctor et al., 2018). Partial glutathione deficiency renders Arabidopsis seedlings highly sensitive to moderately high temperatures, which led to the definition of glutathione as a major player in thermomorphogenesis and heat stress responses in Arabidopsis (Dard et al., 2023). While high temperatures lead to oxidation of the glutathione pool in the cytosol and the nucleoplasm, single null mutants *grxc1*, *grxc2*, *grxc3*, and *grxc4* did not show high-temperature sensitivity. The insensitivity was explained by redundancy between GRX family members. Indeed, functional redundancy has been claimed for GRXC1 and GRXC2 based on the lack of viable double homozygous progeny from self-fertilized *grxc1^−/−^ grxc2^-/+^* plants (Riondet et al., 2012). The developmental stage at which lethality of such mutants is established might be highly informative for identifying key processes or even individual proteins depending on GRX function. However, the stage at which lethality may occur in *grxc1 grxc2* mutants has never been reported and similarly, no information is available on putative redundancy between GRXC3 and GRXC4.

All class I GRXs tested so far are able to reduce glutathionylated substrates (GSSR) or disulfides (RSSR) like bis(2-hydroxyethyl) disulfide (HED) or *E. coli* ribonucleotide reductase with GSH as electron donor (Begas et al., 2017; Couturier et al., 2013; Couturier et al., 2011; Holmgren, 1979). All these established activity assays, however, have the disadvantage that only the reductive half-reaction is considered but not the oxidative reaction. In order to overcome this limitation, an alternative assay based on roGFP2 as a target protein was established and used to analyze the oxidoreductase function of GRXs based on a direct read-out of roGFP2 fluorescence (Begas et al., 2017; A. J. Meyer et al., 2007; Trnka et al., 2020). roGFP2 as a substrate enables bidirectional measurements to analyze the GSH-dependent reduction of oxidized roGFP2 but also the GSSG-dependent oxidation of reduced roGFP2 *in vitro* as well *as in vivo* (Liedgens et al., 2020; Trnka et al., 2020; Zimmermann et al., 2020). In these assays, Arabidopsis GRXC1 has frequently been used as a reference GRX because it efficiently reduces and oxidizes roGFP2 (Begas et al., 2017; Trnka et al., 2020; Zannini et al., 2019). Whether the biochemical separation of GRXC1 and GRXC2 from GRXC3 and GRXC4 observed for the reductive-half reaction with poplar GRXs also occurs for the oxidative half-reaction is not known.

A comprehensive comparison of the common classification systems based on primary and tertiary protein structures failed to correlate the structures to established target specificities. Instead, a functional classification based on electrostatic properties of the respective TRXs and GRXs has been proposed (Gellert et al., 2019). Although endogenous target proteins of GRXs are not known yet, the established interaction with roGFP2 may allow further characterization of Arabidopsis GRXs for their electrostatic properties and the relevance of electrostatics for the respective protein-protein interactions required for oxidoreductase activity. With this it may be possible to further validate the biochemical separation of GRXC1 and GRXC2 from GRXC3 and GRXC4 initially described for poplar (Couturier et al., 2013) and test whether an even more elaborate diversification exists.

Similar to electrostatic compatibility, also the subcellular localization of GRXs is critical for pinpointing their putative target proteins. Class I GRXs are predicted to be targeted to three different subcellular compartments. GRXC5 and GRXS12 contain N-terminal signal peptides that direct both proteins to plastids (Almagro Armenteros et al., 2019). Indeed, GRXC5 and GRXS12 fused with fluorescent proteins showed a clear localization in chloroplasts (Couturier et al., 2011). The localization of the fusion proteins in chloroplasts is also consistent with the presence of GRXC5 and GRXS12 in the chloroplast proteome (Tomizioli et al., 2014). GRXC1 and GRXC2 lack any obvious target signals and thus are generally assumed to be localized in the cytosol as the default compartment. Arabidopsis GRXC1 is predicted to be myristoylated and was indeed found in a set of myristoylated proteins (Majeran et al., 2018; Maurer-Stroh et al., 2002). In addition, GRXC1 was found in a large-scale quantitative proteomics study of plasma membrane-enriched fractions (Elmore et al., 2012). A GFP fusion of poplar GRXC1, however, was found to be distributed equally to the cytosol and the nucleus without labelling membranes when transiently expressed in onion epidermal cells or in *Nicotiana benthamiana* leaves (Riondet et al., 2012; Rouhier et al., 2007). Similarly, GRXC2-GFP has been localized in the cytosol and the nucleus in onion cells and in Arabidopsis suspension culture cells (Riondet et al., 2012; Ströher et al., 2016). Nevertheless, GRXC2 was also found in proteomes of the plasma membrane (Marmagne et al., 2007) and the Golgi (Parsons et al., 2012) raising questions for the exact localization of these proteins. GRXC3 and GRXC4 both contain an N-terminal putative signal peptide that is predicted to direct the proteins to the secretory pathway (Rouhier et al., 2008). Despite the lack of any further known signals for distinct localization or retention in the early secretory pathway, both GRXs have been found in the Golgi proteome (Parsons et al., 2012). In addition, GRXC4 was also found in the ER proteome (Nikolovski et al., 2012). The hydrophobic nature of signal peptides targeting proteins to the ER and of N-terminal transmembrane domains (TMDs) frequently results in erroneous assignments of the respective segments (Nielsen et al., 2019). Based on its redox properties being almost fully reduced in the cytosol and almost fully oxidized in the ER lumen, roGFP2 is a highly suitable binary reporter that self-indicates localization in either compartment (Brach et al., 2009). Expression of N- and C-terminal fusions of the redox-related proteins glutathione peroxidase-like 3 and the ER oxidoreductins 1 and 2 has recently identified all these proteins as type II membrane proteins with one N-terminal TMD (Attacha et al., 2017; Ugalde et al., 2022).

The aim of this work is to provide detailed information on the localization and catalytic properties of the class I GRXs GRXC1-4 in *Arabidopsis thaliana*. For subcellular localization, the proteins were fused with roGFP2 as a self-indicating reporter, to distinguish their putative localization in the cytosol or the secretory pathway when expressed in tobacco (*Nicotiana tabacum*) or Arabidopsis. For functional characterization, recombinant proteins were studied in unidirectional reductive HED assays and also in roGFP2 assays for assessment of bidirectional reducing and oxidizing capacities. The latter assay was complemented by electrostatic modelling and studies of electrostatic compatibility of GRXs with roGFP2. The obtained data reveal a clear separation of the closely related GRXs into two subclasses for both the cytosol and the ER. Knowledge about the localization further allowed the rational generation of double null mutants for both the cytosol and the ER.

## 2 | MATERIALS AND METHODS

### 2.1 | Plant material, mutant selection and growth conditions

Seeds of Arabidopsis (*A. thaliana* [L.] Heyn.) *grxc3-1* (GK-917B02), *grxc3-2* (SALK_134094), *grxc3-3* (SALK_062448C), *grxc4-1* (GK-481F03) and *grxc4-2* (SALK_128264C) T-DNA insertion alleles were obtained from Nottingham Arabidopsis Stock Centre (https://arabidopsis.info/). For genotyping, genomic DNA was extracted from the respective T-DNA mutants according to Edwards et al. (1991). Genotyping was performed by using a left border (LB) primer specific for each mutant line in combination with gene specific primers (P1–10) (*grxc3-1*: LB-Gabi+P1 and P1+P2; *grxc3-2*: LB-Salk+P3 and P3+P4; *grxc3-3*: LB-Salk+P5 and P5+P6; *grxc4-1*: LB-Gabi+P7 and P7+P8; *grxc4-2*: LB-Salk+P9 and P9+P10). See Table S1 for detailed primer information.

The mutants *grxc1* (GABI_509E02), *grxc2* (SALK_076722), and a *grxc1^−/−^ grxc2^-/+^* double mutant were described previously and kindly provided by Jean-Philippe Reichheld (Riondet et al., 2012). Genotyping of these mutants was performed using T-DNA-specific LB primers in combination with gene specific primers for *grxc1* (LB-Gabi+P11 and P11+P12) and *grxc2* (LB-Salk+P13 and P13+P14) (Table S1).

Arabidopsis plants were grown on either Jiffy-7® peat pellets (Jiffy, Zwijndrecht, The Netherlands) or on a soil:sand:vermiculite mixture in the ratio 10:1:1. The soil (Floradur^®^ Multiplication Substrate) was obtained from Floragard (Oldenburg, Germany). Plants were kept in controlled growth chambers under long-day conditions with a diurnal cycle of 16 h light at 22 °C and 8 h dark at 18 °C. The light intensity and relative air humidity were 70 μE m^-2^ s^-1^ and 50 %, respectively. Tobacco (*Nicotiana tabacum* L.) plants were grown in the soil mixture at 22 °C (night/day) under a 16 h photoperiod with a light intensity of 200 μE m^−2^ s^−1^. To grow Arabidopsis seedlings under axenic conditions on agar plates, seeds were surface-sterilized with 70 % (v/v) ethanol, washed three times with dH_2_O, and plated on 0.5× Murashige and Skoog (MS) growth medium (Duchefa Biochemie, Haarlem, The Netherlands) supplemented with 0.1 % (w/v) sucrose, 0.05 % (w/v) MES (pH 5.8, KOH) and 0.8 % (w/v) agar. Plates were incubated vertically under long-day conditions (16 h light at 22 °C and 8 h dark at 18 °C) with a light intensity of 130 μE m^−2^ s^−1^. Seed development was documented by dissecting siliques of self-pollinated plants and counting the number of normal and aborted seeds present in each silique. Siliques were analyzed with a stereomicroscope (Leica M165 FC, Leica Microsystems, Wetzlar, Germany) equipped with a camera (DFC 425 C) and documented using the software LAS V3.8 (Leica Application Suite).

### 2.2 | RNA methods, cloning and site-directed mutagenesis

RNA isolation was performed by using the NucleoSpin^®^ RNA kit (Macherey-Nagel, Düren, Germany) following the manufacturer’s protocol. The RNA was eluted in 50 μL of dH_2_O and quantified with a NanoDrop 2000 spectrophotometer (Thermo Fisher Scientific, Waltham, USA). For reverse transcription of mRNA to cDNA, the RevertAid First Strand cDNA Synthesis Kit (Thermo Fisher Scientific) was used following the manufacturer’s protocol. For determination of transcript abundance in T-DNA-insertional mutants *grxc1* (GABI_509E02), *grxc2* (SALK_076722), *grxc3-1* (GK-917B02), *grxc3-2* (SALK_134094), *grxc3-3* (SALK_062448C), *grxc4-1* (GK-481F03) and *grxc4-2* (SALK_128264C) a semi quantitative RT-PCR was carried out on 1 μL of cDNA with gene-specific primers against *GRXC1-GRXC4* (Table S1 P15-P24).

For subcellular localization, full-length coding DNA sequences (CDS) with or without the stop codons were amplified for GRXC1 (At5g63030), GRXC2.1 (At5g40370.1), GRXC2.2 (At5g40370.2), GRXC3 (At1g77370), GRXC4 (At5g20500), and SEC22 (At1g11890) including attB1 and attB2 flanking sites and cloned into an entry vector using the Gateway^®^ BP clonase II enzyme mix following the manufacturer’s instruction (Invitrogen, Waltham, USA). In addition, CDS versions lacking the N-terminus, including the predicted transmembrane domain (TMD) of GRXC3 (GRXC3_30-130_) or GRXC4 (GRXC4_26-135_) were amplified as well as the N-terminus alone from GRXC2.2 (GRXC2.2_1-43_), GRXC3 (GRXC3_1-29_), and GRXC4 (GRXC4_1-25_). Furthermore, a G_2_A substitution in the putative myristoylation motif of GRXC1 was generated by using primers carrying the respective mutation (Tables S1 and S2). To generate the constructs for the redox-based topology assay (ReTA), roGFP2 was cloned in frame to the C- or N-terminal of the indicated versions of GRXs by using the plasmids pSS01 or pCM01, respectively (Brach et al., 2009; Sparkes et al., 2010). For protein purification, GRXC3_30-130_ was cloned into the plasmid pETG-10A (EMBL Heidelberg, Germany) behind an N-terminal His-tag. GRXC4_26-135_ was cloned into the plasmid pET-28a before a C-terminal His-tag, (Table S2). Cloning of Arabidopsis pET-16b-GRXC1 and pET-16b-GRXC2, pET-30a-roGFP2 was previously described (Brach et al., 2009; A. J. Meyer et al., 2007; Riondet et al., 2012).

### 2.3 | Transient and stable transformation of roGFP2 fusion constructs

Transient transformation of tobacco leaf epidermal cells was performed as described previously (Sparkes et al., 2006) using *Agrobacterium tumefaciens* strain AGL-1 (Lazo et al., 1991) containing the respective binary vectors. Transfected cells were imaged by confocal laser scanning microscopy two days after infiltration. Stable Arabidopsis lines were generated by floral dip according to established protocols (Clough & Bent, 1998) with *A. tumefaciens* (AGL-1) solutions containing 5 % (w/v) sucrose and 0.02 % (v/v) Silwet L-77 as surfactant. Seeds were harvested, germinated on plates and manually screened for fluorescent transformants on a stereomicroscope (Leica M165 FC) equipped with a GFP filter allowing for 470 ± 20 nm excitation and emission at 525 ± 25 nm.

### 2.4 | Confocal laser scanning microscopy and image processing

Leaf disks from infiltrated tobacco plants or five-day-old Arabidopsis seedlings were imaged using a Zeiss LSM 780 confocal laser scanning microscope (CLSM) connected to an Axio Observer.Z1 (Carl Zeiss Microscopy, Jena, Germany) with a 40x lens (C-Apochromat 40x/1.2 W Korr). Images were taken with an additional 6x digital zoom resulting in a pixel size of 90 nm x 90 nm. roGFP2 fluorescence was measured by successive excitation at 405 nm and 488 nm in line switching mode with emission recorded at 505–530 nm for both channels. For colocalization experiments the markers AtWAK2_TP_-mCherry-HDEL (Nelson et al., 2007) was used for the ER and GmManI_1-49_-tdTomato (Hofmann et al., 2009) for the Golgi. mCherry and tdTomato were excited at 543 nm and fluorescence collected between 590–630 nm.

To label the plasma membrane, 7-day-old Arabidopsis seedlings or tobacco leaf pieces were incubated for 10 min in 10 µM FM4-64 (Invitrogen^TM^; T13320; diluted from an 8 mM stock in dimethyl sulfoxide (DMSO). FM-4-64 was excited at 488 nm and fluorescence was collected between 629–665 nm.

Images for Arabidopsis leaves were collected as single images whereas z-stacks with a step width of 0.42 µm were collected for tobacco leaves transiently expressing roGFP2 fusion constructs. Z-stacks were subsequently collapsed as maximum projections to single images and then used for further analysis in ImageJ (https://imagej.nih.gov/ij/). For colocalization with markers for the ER or the Golgi, the respective marker fluorescence was displayed in cyan and roGFP2 fluorescence in magenta for 405 nm excitation and green for 488 nm excitation. To gain information about the relative redox status of roGFP2 in the cytosol and the ER lumen, a scatter plot for the fluorescence intensities after excitation at 405 nm and 488 nm, respectively, was performed using the Colocalization Threshold plugin Coloc2 (https://github.com/fiji/Colocalisation_Analysis/releases). Subsequently, the Pearsońs correlation coefficient above the threshold (R_coloc_) and the gradient (m) of the linear regression fit were determined.

### 2.5 | Heterologous expression in *E. coli* and purification of recombinant proteins

For protein production, the *E. coli* strain BL21 (DE3) was transformed with the recombinant pET-28a, pETG-10A or pET-16b plasmids either with or without the helper plasmid pSBET. Cells were grown at 37 °C to an OD_600_ of ∼0.6 in selective LB medium, and a high level of protein expression was achieved by addition of isopropyl β-D-thiogalactopyranoside (IPTG) to a final concentration of 0.5–1 mM. Regarding GRXC1 and GRXC2, the cultures were harvested after 4 h grown at 37 °C. For GRXC3 and GRXC4, the cultures were harvested after 16 h growth at 21 °C by centrifugation at 5,000 *g* for 15 min and the pellets were re-suspended in 20 mL of Binding Buffer A (20 mM PBS, 500 mM NaCl, 20 mM Imidazole, pH 7.4) with one dissolved tablet of EDTA-free cOmplete (Roche Diagnostics GmbH, Mannheim, Germany). Cell were incubated with 0.5 mM phenylmethylsulfonyl fluoride (PMSF), 1 mg mL^-1^ Lysozyme, and 0.1 mg mL^-1^ DNase for 30 min on ice with constant agitation. Cells were sonicated (4 cycles of 1 min at 30 % power), and the soluble and insoluble fractions were separated by centrifugation for 20 min at 40,000 *g* at 4 °C. The supernatant was immediately transferred to a new 50 mL tube and sterile-filtered through a 0.45 µm filter. The soluble fraction was loaded on a Ni^2+^ affinity column (HisTrap^TM^ Chelating HP Column, GE Healthcare, Uppsala, Sweden) using a peristaltic pump with a flow rate of 1 mL min^-1^. After extensive washing, the protein was eluted from the column using an ÄKTAprime^TM^ plus system (Cytiva, Freiburg, Germany) and a linear 20-500 mM imidazole gradient. The purest fractions as judged by SDS-PAGE (4–20 % polyacrylamide precast gel, BIORAD) gel analysis were pooled and dialyzed against Tris-HCl buffer (30 mM Tris-HCl, pH 8) using PD-10 Desalting Columns (GE Healthcare). The protein concentration was determined by Bradford assays with BSA (bovine serum albumin) as standard. After desalting, the proteins were stored in smaller aliquots at −20°C. GRXC1, GRXC2 and GRXC4 were stored in 30 mM Tris-HCl, pH 8, whereas GRXC3 was stored in 30 mM Tris-HCl, pH 8, 200 mM NaCl to avoid precipitation.

### 2.6 | HED assay for deglutathionylation activity of glutaredoxins

Reduction of 2-hydroxyethyl disulfide (HED) was measured by following NADPH oxidation at 340 nm in the following mixture: 100 mM Tris-HCl, pH 7.9, 1 mM EDTA, 0.2 mM NADPH, 1 mM GSH, 0.6 µg mL^-1^ glutathione reductase (GR; Sigma-Aldrich, St. Louis, USA) and variable concentration of HED (0.1– 5 mM). The concentration of GRX was optimized for all proteins individually (40 nM GRXC1, 40 nM GRXC2, 10 nM GRXC3, and 10 nM GRXC4). The HED assay was performed in a 1 mL cuvette by measuring the absorption every 2 s for 1 min at 340 nm with a NanoDrop 2000 spectrophotometer at 21 °C. The reaction was started by adding GRX after a 3-min pre-incubation time and GRX activity was corrected by subtracting the spontaneous reduction rate observed in the absence of GRX. The activity was expressed as nmol NADPH oxidized s^-1^ nmol^-1^ GRX using a molar extinction coefficient of 6.220 M^−1^ cm^−1^ for NADPH. Three independent reactions were performed at each concentration, and apparent *k*_cat_ and *K*_m_ values were calculated by nonlinear regression using the Michaelis–Menten equation.

### 2.7 | roGFP2 assay for oxidoreductase activity of glutaredoxins

Interaction of GRXs with roGFP2 was analyzed *in vitro* by ratiometric time-course measurements on a fluorescence plate reader (CLARIOstar, BMG Labtech, Ortenberg, Germany) with excitation at 400 ± 5 nm and 480 ± 5 nm and detection of emitted light at 520 ± 5 nm. The reaction mixture contained 0.1 M potassium phosphate buffer with 1 mM EDTA, pH 7.4 and 1 µM roGFP2 with the respective GRXs. To fully reduce roGFP2, 10 mM DTT was added for at least 30 min and then removed by gel filtration (Zebra Spin Desalting Columns, Thermo Fisher Scientific). To test the capability of GRXs to catalyze the reduction of roGFP2, GSH was added to a final concentration of 2 mM together with 10 U yeast GR (Sigma-Aldrich) and 100 µM NADPH. To test GRX-dependent oxidation of roGFP2, 40 µM GSSG was added to the wells. For determination of the dynamic range of the sensor and adapting the gain values for both fluorescence channels roGFP2 was either fully oxidized or full reduced through incubation with either 10 mM H_2_O_2_ or 10 mM DTT, respectively. A basal background fluorescence of the buffer was subtracted from fluorescence reads for all samples. The degree of roGFP2 oxidation was analyzed as described before (Aller et al., 2013). The respective turnover rate *k*_cat_ was calculated as described previously (Trnka et al., 2020).

### 2.8 | Phylogenetic analysis

The Arabidopsis GRXC1 sequence was used to identify protein orthologues in *Arabidopsis thaliana*, *Hordeum vulgare*, *Medicago truncatula*, *Oryza sativa*, *Populus trichocarpa*, *Solanum lycopersicum*, *Solanum tuberosum*, *Vitits vinifera*. Sequences were retrieved from version 13 of the Phytozome platform (Goodstein et al., 2012) (https://phytozome-next.jgi.doe.gov/) through a pblast search. The corresponding gene identifiers are listed in Table S3. Multiple sequence alignments were built with SeaView (Gouy et al., 2010) using Clustal Omega with default settings. To generate the phylogenetic tree with SeaView (PhyML, default settings: BioNJ starting tree, LG model, NNI improvement), sequences were trimmed at their termini so that all sequences spanned residues between helix α0 and helix α4. 98 (Figure 1) or 108 (Figure S1) positions were used for tree-building.

**FIGURE 1.**
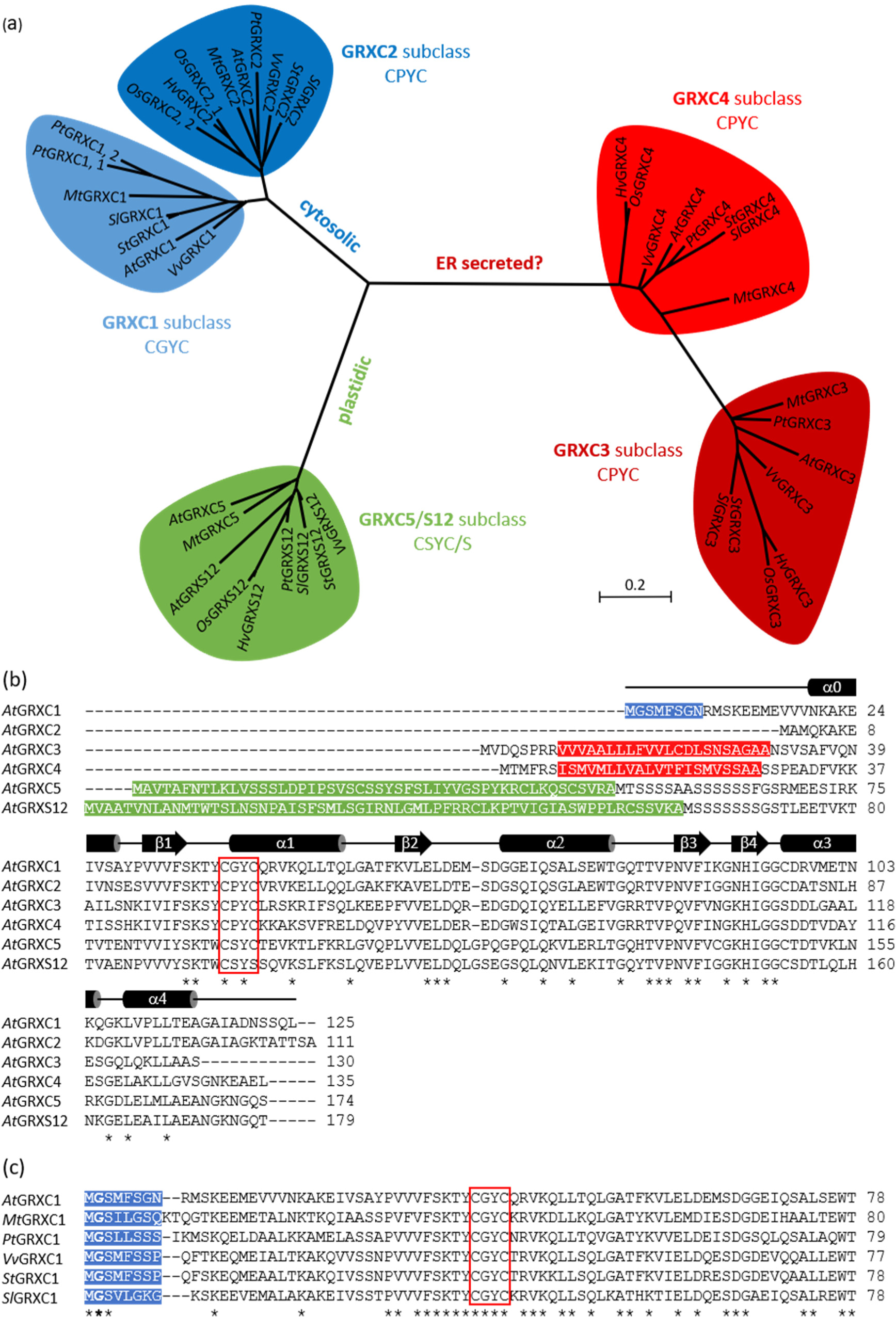
Phylogenetic analysis of representative class I glutaredoxins from spermatophytes. (a) Unrooted tree of class I glutaredoxins. The scale bar represents 0.2 substitutions per amino acid position. Species names are abbreviated: *At* = *Arabidopsis thaliana*; *Hv* = *Hordeum vulgare*; *Mt* = *Medicago truncatula*; *Os* = *Oryza sativa*; *Pt* = *Populus trichocarpa*; *Sl* = *Solanum lycopersicum*; *St* = *Solanum tuberosum*; *Vv* = *Vitits vinifera* and GRXC1 appears only with the evolution of dicotyledonous plants (Couturier et al., 2009). (b) Sequence alignment of class I glutaredoxins from Arabidopsis. The red box indicates the C(P/G/S)Y(C/S) active site motif. A putative myristoylation motif at the N-terminus of GRXC1 is highlighted in blue. Putative transmembrane domains predicted by the PRED-TMR algorithm are highlighted in red. Putative plastid-targeting sequences of GRXC5 and GRXS12 are highlighted in green. The nomenclature of secondary structure elements corresponds to the classical thioredoxin fold, which starts with β1 (Martin, 1995). Fully conserved positions are marked with an asterisk. (c) Sequence alignment of different plant GRXC1 paralogs. The red frame indicates the four amino acids comprising the active site motif; putative motifs for myristoylation of a conserved N-terminal glycine residue in second position (bold), as predicted by the NMT Predictor, are highlighted in blue. Fully conserved positions are marked with an asterisk.

### 2.9 | Molecular modeling and electrostatic calculations

The structures of AtGrxC1, C2, C3, C4 were modelled using SWISS-MODEL (Waterhouse et al., 2018). The modelled structures were subjected to short all-atomistic molecular dynamics (MD) simulations. Gromacs 2021.3 (Van Der Spoel et al., 2005) with AMBER-99ff-ILDN (Sansenya et al., 2010) force field were used for these MD simulations. The proteins were put in a cubic box with 1 nm distance between protein and the edges of the box, and solvated with TIP3P (transferable intermolecular potential with 3 points) water. The simulation box was neutralized with Na^+^ and Cl^-^ ions. The energy of the system was minimized with steepest descent algorithm until the systems had reached 1000 kJ·mol^-1^·nm^-1^. Next, the system was equilibrated for 100 ps under NVT conditions (*i.e.*, constant number of molecules, volume, and temperature of 300 K) and for 100 ps under NPT conditions (*i.e.*, constant number of molecules, pressure of 1 bar, and temperature of 300 K). Afterwards, the simulations with duration of 50 ns were performed. The last frames of the simulations were used for the calculation of electrostatic potentials. For that, we used the method described in (Gellert et al., 2019). In brief, the pdb2pqr (Dolinsky et al., 2007) tool was used for the assignment of atomic charges and atomic radii to the atoms of the structures with the Amber99ff force field. The Adaptive Poisson-Boltzmann Solver (APBS) (Baker et al., 2001) tool of visual molecular dynamics (VMD) (Humphrey et al., 1996) was used for the calculation of the electrostatic potentials with the parameters described in (Gellert et al., 2019). The hierarchical clustering was performed using pairwise calculation of earth-mover’s distances (EMDs) for four randomly selected points on the surface, repeated 10 times. The EMDs were then used for the unweighted pair group method with arithmetic mean (UPGMA) for creation of the clustering tree.

### 2.10 | Statistical analysis

For the statistical analysis of the catalytic efficiency of GRXC1-4 (*k_cat_*/*K_m_*) a one-way ANOVA with a comparison between the mean values of each column was performed in GraphPad Prism (version 8.3) All biochemical experiments were performed with each three biological and three technical replicates. Data were calculated as means with standard deviation. Statistically significant differences between samples were determined using one-way ANOVA analyses followed by a Holm-Sidak test in GraphPad Prism 8.3 (P > 0.05: ns; P ≤ 0.05: *P ≤ 0.01: **P ≤ 0.001: ***).

## 3 | **RESULTS**

### 3.1 | Sequence analysis and targeting predictions for class I glutaredoxins

Phylogenetic analysis of class I GRXs from several spermatophytes based on the core TRX fold (β1, α1, β2, α2, β3, β4, α3, α4) revealed that the proteins group into three larger clusters of which two can be further subdivided (Figure 1a). Based on assignment of all six class I GRXs encoded in the Arabidopsis genome, the five subclasses C1, C2, C3, C4, and C5/S12 can be separated, which are annotated as cytosolic, plastidic and secreted proteins. Sequence analysis of class I GRXs from Arabidopsis revealed distinct differences particular at their N-termini (Figure 1b). GRXC5 and GRXS12 contain a plastid target peptide (Figure 1b, green letters; Figure S1) and have been experimentally confirmed as plastid proteins (Couturier et al., 2011). Therefore, both proteins were excluded from further analysis in this study. The other two larger clusters both separate into two subclasses, which in Arabidopsis contain one GRX isoform each (Figure 1a; Figure S1). Even when the phylogenetic analysis is restricted to the core GRX domain without any N-terminal extensions, the subclasses C1 and C2 form one cluster that clearly separates from the cluster with subclasses C3 and C4. Both, GRXC3 and GRXC4, contain hydrophobic domains spanning 21–22 amino acids in their N-terminal regions for which the consensus prediction based on different bioinformatics algorithms is a cleavable N-terminal target peptide for the secretory pathway (Figure 1b, white letters on red background; Figure S2). Individual algorithms, however, differ in interpretations of the N-termini of GRXC3 and GRXC4 and predict either cleavable targeting peptides, lack of any targeting information of hydrophobic domains, or the presence of TMDs with orientation of the N-termini either inside or outside the cytosol. In case of a cleavable signal, this would lead to a soluble protein that in the absence of any retrieval signal would be destined for secretion. Conversely, the presence of a TMD would possibly anchor the protein to membranes along the secretory pathway or the plasma membrane with a distinct orientation of the catalytic domain to one or the other side of the membrane. In the latter case, the orientation predictions of the N-terminus of GRXC4 are not unanimous, which highlights the need for experimental validation of protein localization and their topology if they are anchored to a membrane. GRXC1 and the annotated spliced variant for GRXC2, GRXC2.1, do not contain any predicted target peptide for organelle targeting, which suggests default cytosolic localization. A second annotated splice variant, GRXC2.2, however, contains a longer N-terminus with a 21-amino-acid hydrophobic domain that may target the protein to the endomembrane system (ARAMEMNON, Schwacke et al. (2003); SUBA, Hooper et al. (2017)). Notably, the predicted hydrophobic domain in this case replaces the β_1_-sheet of the canonical TRX fold motif, which would most likely disturb the normal catalytic GRX function (Figure 1b). While GRXC1 does not contain any cleavable target peptide, the sequence does contain a putative N-terminal myristoylation motif (Figure 1b, white letters on blue background). Analysis of the amino acid sequences of GRXC1 orthologues from other spermatophytes shows a high degree of similarity of the typical (M)GxxxS/Txx myristoylation motif (Turnbull & Hemsley, 2017) although the motif is not fully conserved in all cases (Figure 1c). GRXC1 and its orthologues are thus predicted as cytosolic proteins with localization at membranes.

### 3.2 | roGFP2 self-indicates localization of proteins in the ER or in the cytosol

To test whether GRXC1-4 are localized in the cytosol or the secretory pathway, we exploited the steep *E*_GSH_ gradient between the cytosol and the ER lumen by using the fluorescent *E*_GSH_ sensor roGFP2 as a self-reporting tag. Since roGFP2 will be almost fully reduced when facing the cytosol and almost fully oxidized when facing the lumen, it provides a binary ratiometric readout depending on its orientation relative to the ER membrane (Figure S3) (Brach et al., 2009). Colocalization analysis of the fluorescence intensities at both excitation channels for each pixel in a scatter plot provides an even more straightforward analysis showing a much higher slope of the regression line for the cytosol (m = 2.38) than for the ER lumen (m = 0.43) for soluble proteins expressed in Arabidopsis (Figure S3a,b). Similar results with clear separation of luminal and cytosolic roGFP2 were observed for the single spanning SNARE SEC22 tagged at either end with roGFP2 and transiently expressed in tobacco (*Nicotiana tabacum*) leaves (Figure S3c,d). Given that the readout is binary, no further quantification of the fluorescence is necessary and even the simple merge of the two fluorescence channels resulting in a light green merge color for the cytosol and light magenta for the ER lumen is fully informative.

### 3.3 | GRXC1 and GRXC2 are cytosolic proteins

To prove the subcellular localization of the different GRXs and, in case they are anchored to endomembranes or the plasma membrane, also their orientation relative to the membrane, we fused roGFP2 to either the N- or the C-termini of GRXC1-4. The roGFP2 fusions to GRXC1 and both annotated splice variants of GRXC2 (GRXC2.1 and GRXC2.2) were transiently expressed in tobacco leaves and stably expressed in Arabidopsis. For all these construct permutations, the merge of both roGFP2 fluorescence channels was consistently dominated by the 488 nm channel and thus showed a green merge color (Figure 2; Figure S4). Based on comparison with control constructs for the cytosol and the ER (Figure S3), this unequivocally indicates all respective protein fusions to be localized in the cytosol. While roGFP2-GRXC1 and roGFP2 fusions to both termini of both GRXC2 splice variants were found to share a cytosolic-nuclear localization with even fluorescence distribution, the localization of GRXC1-roGFP2 was different. In pavement cells of tobacco leaves, there was some cytosolic and nuclear signal detectable, but especially the nuclear signal was far less intense than with all other fusion proteins (Figure S4b). In addition, GRXC1-roGFP2 fluorescence colocalized with ER-targeted mCherry at the nuclear envelope and the ER, and resulted in fluorescently labelled punctate structures that were not seen for any other construct (Figure S4b). In stably transformed Arabidopsis lines, the fluorescence was mainly confined to the outer rim of the cells and even more pronounced punctate structures were present (Figure 2b). Strong transient co-expression of GRXC1-roGFP2 with the nominal Golgi marker GmMan11-49-tdTomato in tobacco resulted in colocalization at the nuclear envelope and the Golgi (Figure 3a). In addition, GRXC1-roGFP2 partially localized to the outer rim of the cells where it colocalized with the plasma membrane dye FM4-64 in tobacco and Arabidopsis (Figure 3c,d). This suggests that GRXC1-roGFP2 harboring compartments are most likely part of the endomembrane system and the plasma membrane. To test whether the observed localization of GRXC1-roGFP2 was dependent on myristoylation, we mutated the putative myristoylation site by replacing glycine in position 2 with alanine. Expression of GRXC1_(G2A)_-roGFP2 in tobacco resulted in an even cytosolic-nuclear distribution of fluorescence without labeling of membranes (Figure 3b). Consistently, these results show that GRXC1 and GRXC2 are both cytosolic with GRXC1 being attached to membranes through N-terminal myristoylation.

**FIGURE 2.**
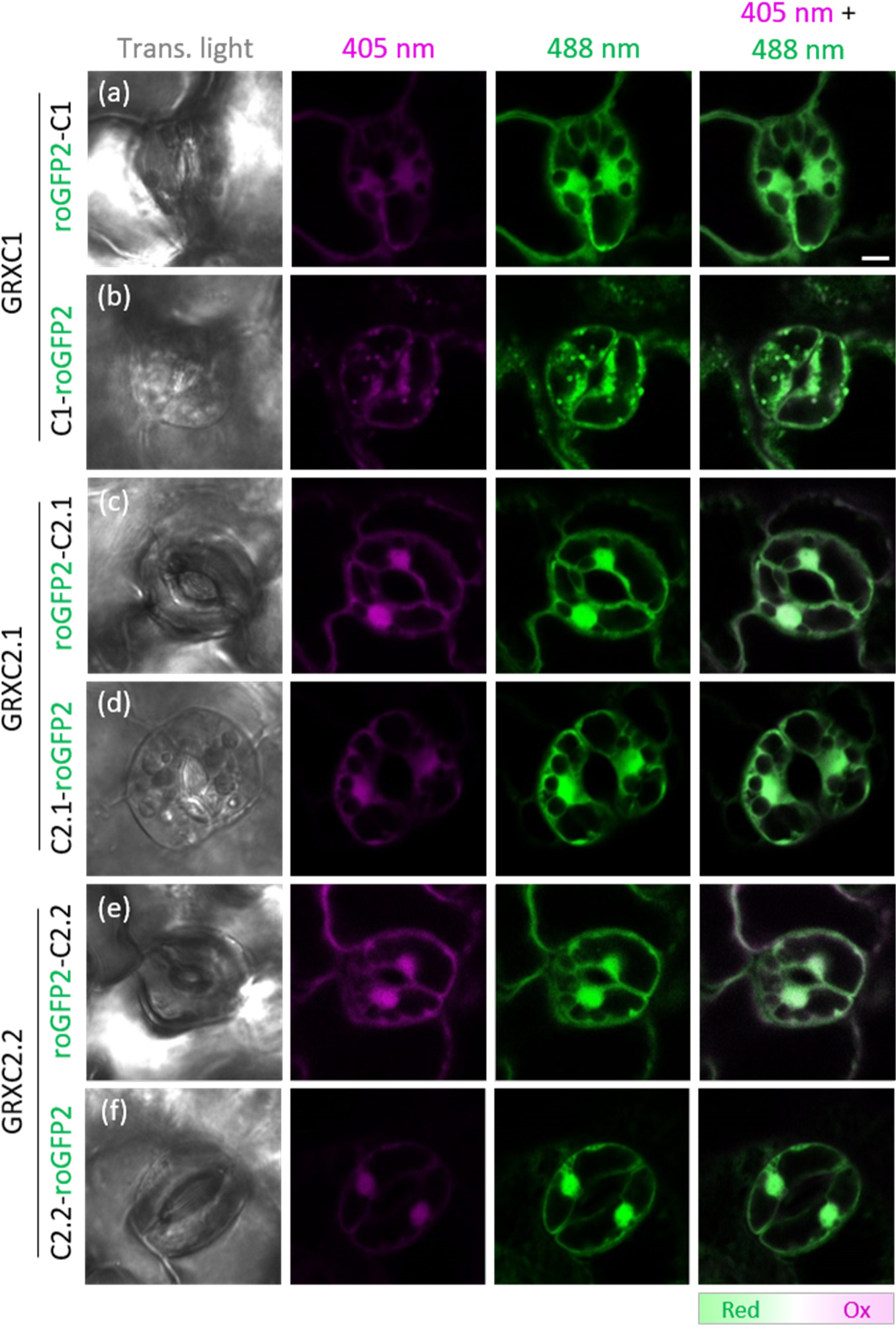
Subcellular localization of GRXC1, GRXC2.1 and GRXC2.2 in *Arabidopsis*. (a-f) Representative confocal microscopy images of guard cells from 7-day-old seedlings stably expressing roGFP2 fusions to the N- and C-termini of GRXC1 (a,b), GRXC2.1 (c,d) and GRXC2.2 (e,f). Images show roGFP2 fluorescence collected at 505–530 nm after excitation with either 405 nm (magenta) or 488 nm (green). The green color obtained after merging these two channels indicate roGFP2 for all constructs to be reduced. Bar = 5 µm.

**FIGURE 3.**
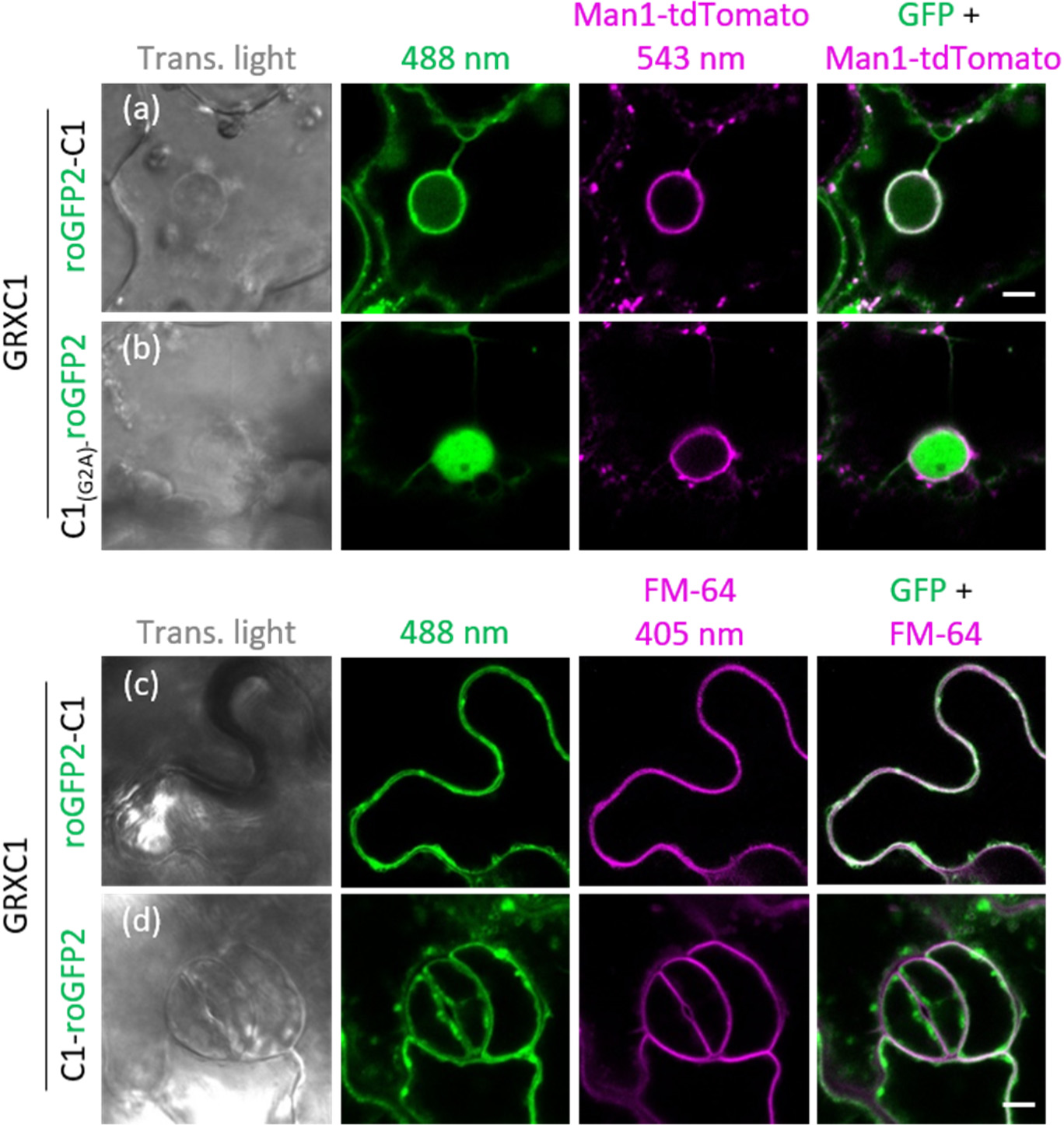
GRXC1 is membrane-anchored through N-terminal myristoylation. (a,b) Representative confocal microscopy images of pavement cells in the leaf epidermis of tobacco transiently expressing roGFP2 fusions to C-termini of GRXC1 (a), and the mutated variant GRXC1_(G2A)_ (b). Images show roGFP2 fluorescence collected at 505–530 nm after excitation with 488 nm (green). To test for colocalization along the secretory pathway, leaves were co-infiltrated with the nominal Golgi marker GmManI_1-49_-tdTomato, which in strong transient overexpression results in labeling of the ER and the Golgi. Fluorescence of tdTomato was excited at 543 nm and collected at 590-630 nm (magenta). The merge of the 488 nm and 543 nm channels indicates colocalization by a light green to white color. Bar = 5 µm. (c,d) Representative confocal images of pavement cells transiently expressing GRXC1-roGFP2 in tobacco leaves (c) or in stably transformed Arabidopsis lines (d). In both cases, leaf pieces were additionally stained with FM4-64.

### 3.4 | GRXC3 and GRXC4 are type II membrane proteins in the endomembrane system

To experimentally validate the prediction for GRXC3 and GRXC4 as being targeted to the secretory pathway and to test whether the proteins are secreted or kept in the endomembrane system, we again exploited the redox-sensitive features of roGFP2 and expressed the respective N- and C-terminal fusion proteins in leaves of tobacco and Arabidopsis. With the exception of GRXC3-roGFP2 for which despite several attempts no stable Arabidopsis reporter line could be recovered, all construct permutations resulted in a fluorescence pattern indicating endomembrane labeling (Figure 4, Figure S5). In addition, the C-terminal fusions of GRXC3 and GRXC4 with roGFP2 resulted in net-like fluorescence distribution, which was particularly evident in pavement cells of tobacco leaves (Figure S5). Co-expression with the ER-marker AtWAK2_TP_-mCherry-HDEL in tobacco leaves shows GRXC3 and GRXC4 to be ER-targeted (Figure S5). The merge of the two roGFP2 images obtained after sequential excitation at 405 nm and 488 nm, consistently showed green for roGFP2 fused to the N-termini of GRXC3 and GRXC4 and magenta when fused to the C-termini (Figure 4; Figure S5).

**FIGURE 4.**
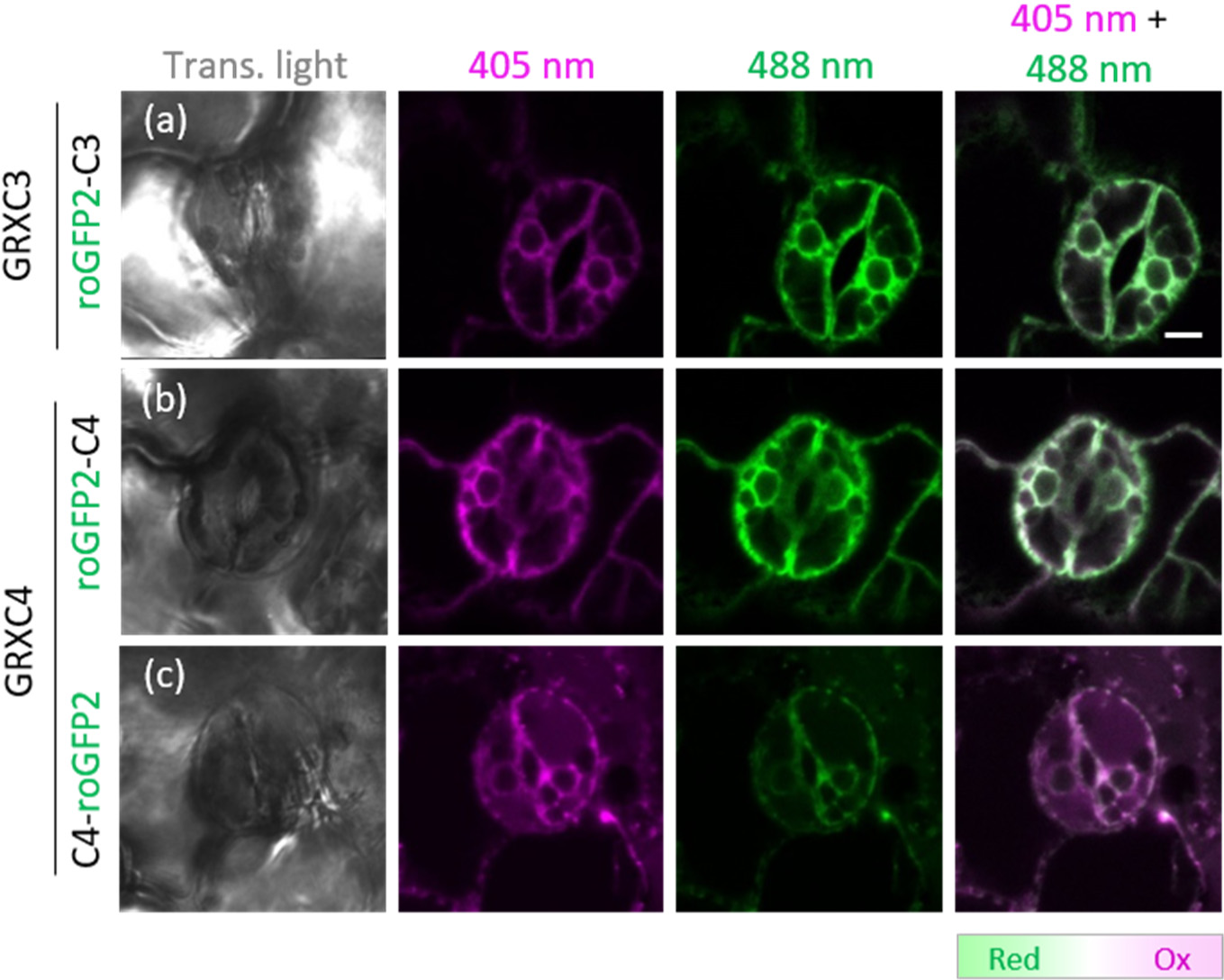
Subcellular localization of GRXC3 and GRXC4 in *Arabidopsis*. (a-c) Representative confocal microscopy images of guard cells from 7-day-old seedlings stably expressing roGFP2-GRXC3 (a) or GRXC4 with roGFP2 fused to the N- or the C-termini (b,c). Despite several attempts, no stable lines with expression of GRXC3-roGFP2 could be recovered. Images show roGFP2 fluorescence collected at 505–530 nm after excitation with either 405 nm (magenta) or 488 nm (green). The colors obtained after merging these two channels indicate the oxidation level of roGFP2 on a scale from green for reduced protein to magenta for oxidized protein. Bar = 5 µm.

To further test whether the putative N-terminal TMDs of GRXC3 and GRXC4 alone were sufficient to mediate and maintain their subcellular localization, we fused roGFP2 to either the N- or C-termini of the predicted TMDs of GRXC3 and GRXC4 and expressed these constructs in tobacco. In all combinations, the roGFP2 fluorescence excited at 405 nm colocalized with ER-targeted mCherry (Figure 5) suggesting that the TMDs are sufficient for ER targeting of GRXC3 and GRXC4. It was noted though that the fusion of roGFP2 to the N-terminus of the GRXC3 TMD remained at least partially in the cytosol and the nucleoplasm, which may indicate partial masking of the respective targeting signal (Figure 5a). Expression of the N-terminal fusions roGFP2-GRXC3_1-29_ and roGFP2-GRXC4_1-25_ resulted in higher fluorescence when excited at 488 nm than at 405 nm, while the opposite was seen with the C-terminal fusions GRXC3_1-29_-roGFP2 and GRXC4_1-25_-roGFP2 (Figure 5). The binary readout for the merge images with N-terminal fusions in green and C-terminal fusions in magenta further support the notion that GRXC3 and GRXC4 are type II proteins in the ER membrane.

**FIGURE 5.**
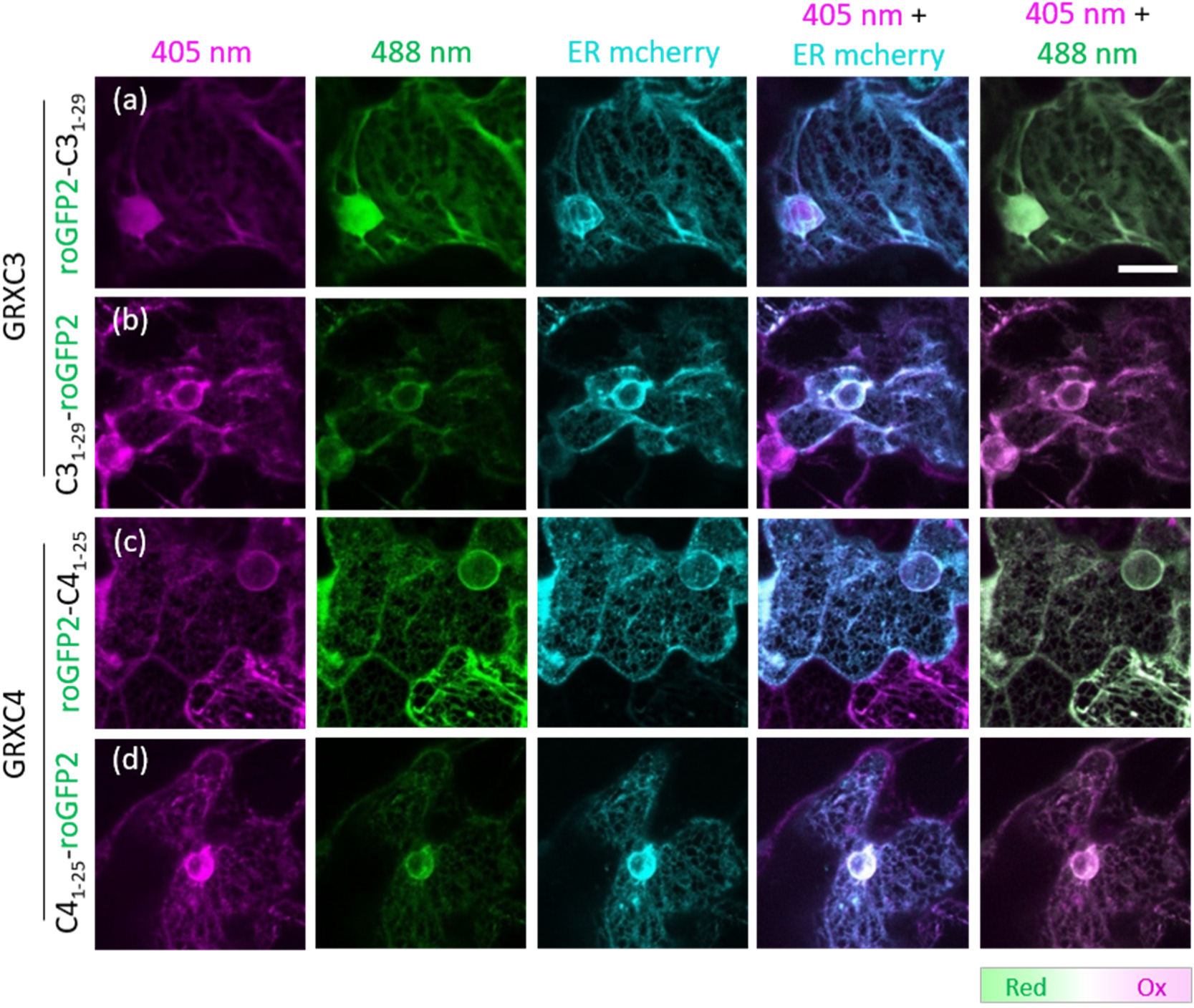
Subcellular localization of the transmembrane domains of GRXC3 and GRXC4 in tobacco. (a-d) Representative confocal microscopy images of pavement cells in the leaf epidermis transiently expressing roGFP2 fusions to the N- and C-termini of GRXC3_1-29_ (a,b) or GRXC4_1-25_ (c,d). Images show roGFP2 fluorescence collected at 505–530 nm after excitation with either 405 nm (magenta) or 488 nm (green). For ER colocalization, leaves were co-infiltrated with the marker AtWAK2_TP_-mCherry-HDEL (cyan). The colors obtained by merging the 405 nm and 488 nm roGFP2 channels indicates the oxidation level of roGFP2 on a scale from green for reduced protein to magenta for oxidized protein. Bar = 20 µm.

When GRXC3 and GRXC4 where expressed without their TMDs, the fusion proteins were found as soluble cytosolic and nuclear proteins that were excluded from the ER (Figure S6).

Altogether, these results suggest that GRXC3 and GRXC4 are type II ER-membrane proteins with their N-termini oriented towards the cytosol and their C-termini including the GRX domain towards the ER-lumen. Both proteins are anchored to the membrane by a single-spanning TMD of ∼21 amino acids at their N-termini.

### 3.5 | The double mutants *grxc1 grxc2* and *grxc3 grxc4* lack obvious phenotypes

Pronounced sequence similarities between GRXC1 and GRXC2 as well as GRXC3 and GRXC4 in combination with the subcellular localization of these two pairs of closely related GRXC isoforms suggests that the respective proteins may have at least partially redundant functions. Based on the expectation that higher order null mutants would possibly cause distinct phenotypes and thus provide insight into the physiological function of class I GRXs, we isolated null mutants for all four GRXs and generated the respective double mutants. The hypothesis for functional redundancy between GRXC1 and GRXC2 was fostered by earlier work in which a line homozygous *grxc1* and heterozygous *grxc2*/*GRXC2* that carried a *grxc1*-*grxc2* recombinant chromosome was isolated but did not give rise to double homozygous *grxc1 grxc2* mutants (Riondet et al., 2012). Because Riondet and colleagues did not specifically report at which developmental stage lethality may occur, we re-investigated the same mutants carrying T-DNA insertions in the first intron (*grxc1*) and the second intron (*grxc2*), respectively. Consistent with the original report, no residual transcripts were found in the two lines confirming both lines as null mutants (Figure S7a,b). In our hands, however, fully viable double-homozygous *grxc1 grxc2* mutants could be isolated (Figure S7). Since it was previously reported that siliques of *grxc1^−/−^ grxc2^-/+^* plants lacked several seeds, we further analyzed the siliques of the *grxc1 grxc2* double mutant in comparison with siliques of wild-type plants. Here, no abortion of seeds nor any hints for defects in fertilization were found (Figure S7c). Based on this observation, the *grxc1 grxc2* double mutant together with the respective single mutants were further investigated for their germination rate, root length during seedling stage and vegetative growth. For none of these parameters pronounced phenotypic characteristics deviating from the wild type could be identified (Figure S7d-f). To enable functional analysis for GRXC3 and GRXC4, several T-DNA lines for both loci were selected (Figure S8a,b). For all alleles, homozygous mutants were isolated and validated by PCR. Transcript analysis showed no loss of GRXC3 transcript for *grxc3-2*, which carries an insertion in the 5’-UTR. For *grxc3-1*, which contains an insertion in the third intron, the primer pair P19+P20 gave rise to a weak band indicating the presence of residual transcript (Figure S8c). Using an exon-exon spanning reverse primer (P21) annealing to parts of exon two and exon three in the coding region of GRXC3, a similar transcript level as in the wild type was detected. Because this may still give rise to a truncated protein, a third T-DNA line named *grxc3-3* with an insertion in the first exon was isolated. In this case, a complete loss of transcript was confirmed with primer combinations P19+P20 and P19+P21 (Figure S8c). For *GRXC4,* the allele *grxc4-1* carrying an insertion in the promoter region had transcript levels similar to the wild type control. In *grxc4-2* plants carrying a T-DNA at +275 bp relative to the start codon, a complete loss of *GRXC4* transcript was observed (Figure S8d). Because neither *grxc3-1*, *grxc3-3* nor *grxc4-2* showed any obvious phenotype, we crossed the respective lines to test for functional redundancy between GRXC3 and GRXC4. Double homozygous mutants could be isolated but did not show any obvious phenotype under normal growth conditions (Figure S8e-h).

### 3.6 |Class I glutaredoxins in the cytosol and the ER differ in their oxidoreductase activities

In the absence of clear phenotypes, we next asked whether the evolutionary separation of subclasses and the localization of GRXC1 and GRXC2 in the cytosol with an *E*_GSH_ of −315 mV (A. J. Meyer et al., 2007) and GRXC3 and GRXC4 in the ER lumen with an *E*_GSH_ of −240 mV (Ugalde et al., 2022) may correlate with biochemical properties of the respective oxidoreductases. Class I GRXs are generally characterized as oxidoreductases capable of mediating glutathionylation and deglutathionylation of target proteins (Ströher & Millar, 2012). We first tested their reductase activity using the classical HED assay, which measures the reduction of glutathionylated β-mercaptoethanol (β-ME-S-SG) (Figure S9a) (Begas et al., 2015). In all cases, a hyperbolic relationship between the reaction rate and the concentration of HED was obtained (Figure S9b-e). The apparent *K*_m_ values for the cytosolic GRXC1 and GRXC2 were 170 µM and 397 µM, respectively (Table S4). In contrast, for GRXC3 and GRXC4 we found higher *K*_m_ values of 809 µM and 1127 µM (Table S4). At the same time, turnover rates *k*_cat_ for GRXC3 (131 s^-1^) and GRXC4 (219 s^-1^) were higher than for GRXC1 and GRXC2 for which *k*_cat_ values of 17 s^-1^ and 32 s^-1^ were determined. Further analysis resulted in higher catalytic efficiencies (*k*_cat_*/K*_m_) for GRXC3 and GRXC4 than for GRXC1 and GRXC2 (Figure 6a; Table S4).

**Figure 6.**
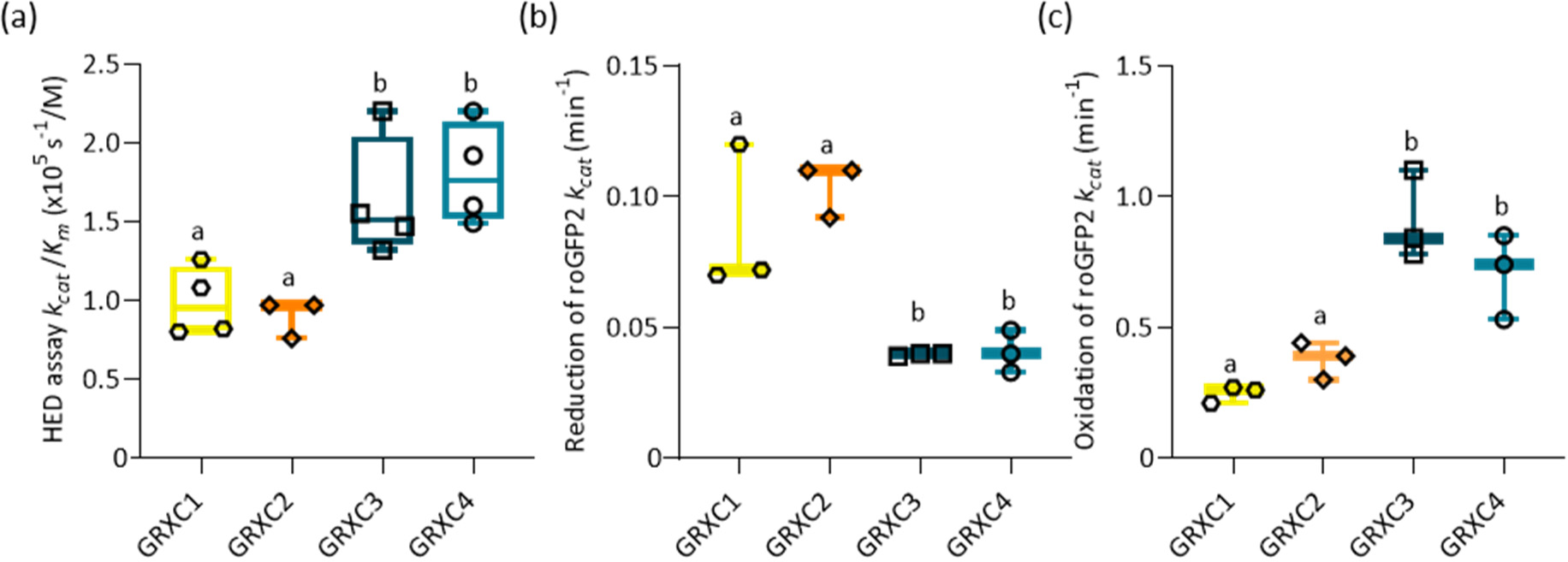
Comparison of catalytic activities of cytosolic and luminal class I GRXs. (a) Catalytic efficiency (*k*_cat_/*K*_m_) of GRXC1-C4 in the HED assay, from three to four biological replicates (*n*=3-4). (b-c) Catalytic activity apparent (*k*_cat_^app^) of GRXC1-C4 determined in bidirectional roGPF2 oxidoreductase activity assay. *n*=3. The box plots show the interquartile range between lower and upper quartiles (Box) with the median (line in the center) and the min and maximum values (whiskers). Statistical analyses were performed using ANOVA with Holm-Sidak test. Different letters indicate statistically different groups (P<0.05).

Since the HED assay allows only unidirectional analysis of reductive capacities, we next exploited roGFP2 as a GRX-target protein for which reversible thiol-disulfide reactions can be directly observed by monitoring the chromophore fluorescence in real time (A. J. Meyer et al., 2007; Trnka et al., 2020). In the presence of GR and NADPH to maintain maximum reduction of the glutathione pool, GRXC1 and GRXC2 efficiently catalyzed the reduction of oxidized roGFP2 with 2 mM GSH with turnover rates of 0.09 min^-1^ and 0.1 min^-1^ (Figure 7a; Table S4; Figure S10). Consistent with results from the HED assay, GRXC3 or GRXC4 were significantly less efficient in reducing roGFP2 with turnover rates of 0.04 min^-1^, respectively (Figure 6b; Figure 7a; Table S4; Figure S10). For the oxidation of pre-reduced roGFP2 with 40 µM GSSG, however, the reaction was much faster in the presence of GRXC3 or GRXC4 with significant different turnover rates of 0.9-0.71 min^-1^ than with GRXC1 or GRXC2, for which *k*_cat_ values were between 0.25 and 0.38 min^-1^ (Figure 6c; Figure 7b; Table S4; Figure S11). While full oxidation of 1 µM roGFP2 with 40 µM GSSG was achieved within 3 minutes by GRX3 and GRXC4, the reaction catalyzed by GRXC1 and GRXC2 took ∼12 min to reach full oxidation of roGFP2 (Figure 7b). This result indicates that GRXC3 and GRXC4 have an enhanced capacity to oxidize target proteins.

**FIGURE 7.**
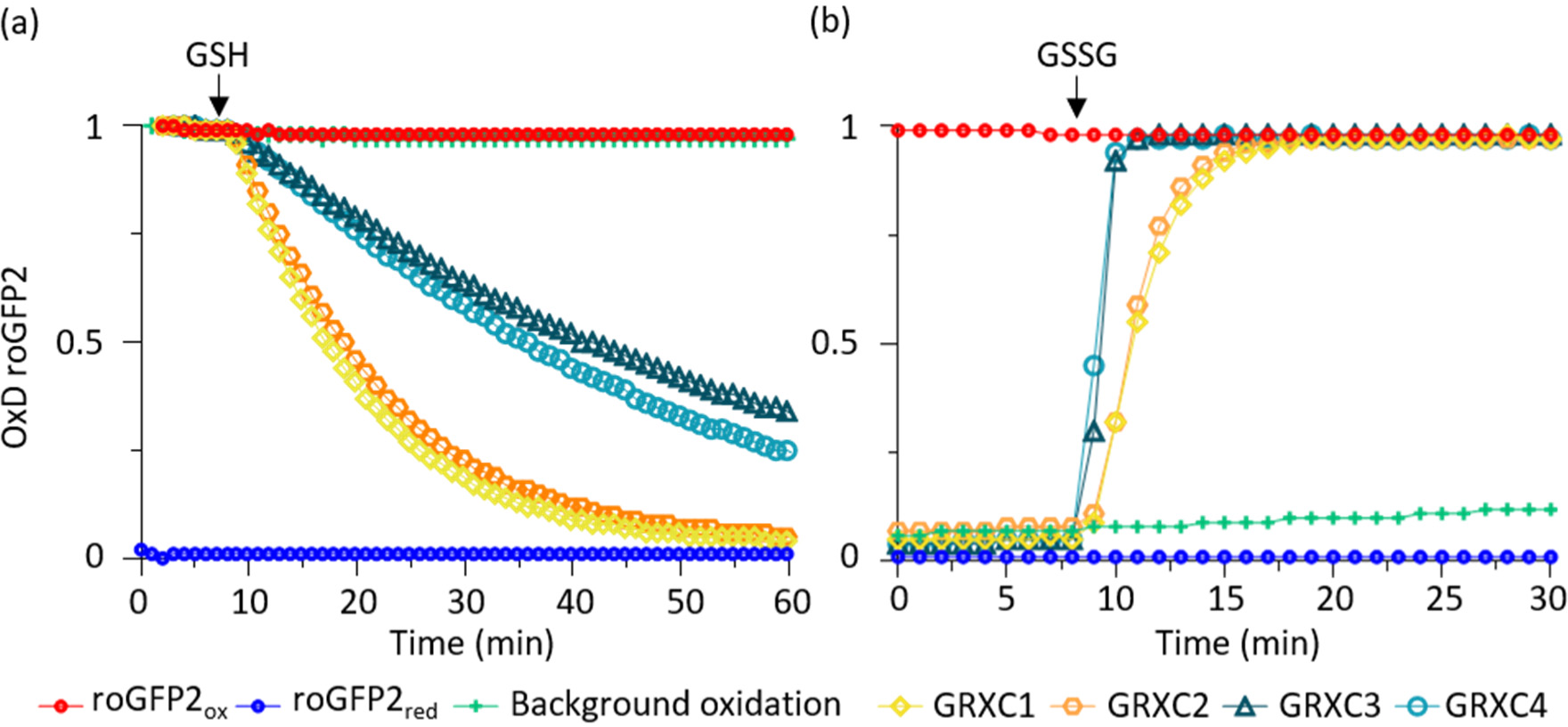
Oxidoreductase activity of class I GRXs in roGFP2 interaction assays. (a,b) GSH-dependent reduction of 1 μM oxidized roGFP2 (a) and GSSG-dependent oxidation of 1 μM reduced roGFP2 (b) in presence of 1 µM GRXs. The arrow indicates the injection of 2 mM GSH (a) or 40 µM GSSG (b). Full reduction or oxidation was determined by incubating roGFP2 with 10 mM DTT or 10 mM H_2_O_2_, respectively. All data points are means from three technical replicates.

Differences in catalytic properties of enzymes are generally explained by structural alterations in the active site. Here, however, comparison of the active sites of GRXC1-4 does not highlight any change that would explain the catalytic separation of two subclasses observed in biochemical assays. An amino acid substitution that correlates with the functional separation is the presence of a third cysteine residue in GRXC1 and GRXC2 at the start of the helix α3, which might be situated close to the catalytic motif on helix α1 (Figure S1). In GRXC3 and GRXC4, this cysteine is substituted by serine. Structural modelling, however, suggests that the distance between the C-terminal cysteine and the closest active site cysteine is 9.4 Å and thus too far away for formation of an internal disulfide bridge. Indeed, Couturier et al. (2013) showed that mutation of this cysteine residue does not impact on the catalytic properties of the poplar protein in the HED assay. Multiple other residues, especially on the helix α2 opposing the active site motif have been highlighted as candidates with some potential impact of the catalytic activity (Figure S1; Liedgens et al., 2017). Detailed analysis of the respective complexity of these amino acid patterns would be far beyond the scope of this work.

In an alternative model, substrate specificity and catalytic efficiency of TRXs and GRXs has been explained by the extent of a negative electric field of the redoxins reaching into the solvent outside the active site, and electrostatic and geometric complementary contact surfaces (Berndt et al., 2015; Gellert et al., 2019). To further test the hypothesis that catalytic activities of GRXs are based on electrostatic compatibilities with their substrates, we modelled the electrostatic surface of all four GRXs and roGFP2 as the respective target protein in our test system. In the electrostatic similarity tree, the four GRXs were clearly separated and formed two clusters with GRXC1 and GRXC2 at one end and GRXC3 and GRXC4 at the other (Figure 8). Most notably, to achieve electrostatic compatibility with the roGFP2 substrate, the two subclasses must adopt different orientations towards the substrate, *i.e.* rotated by 180° (see Figure 8b).

**FIGURE 8.**
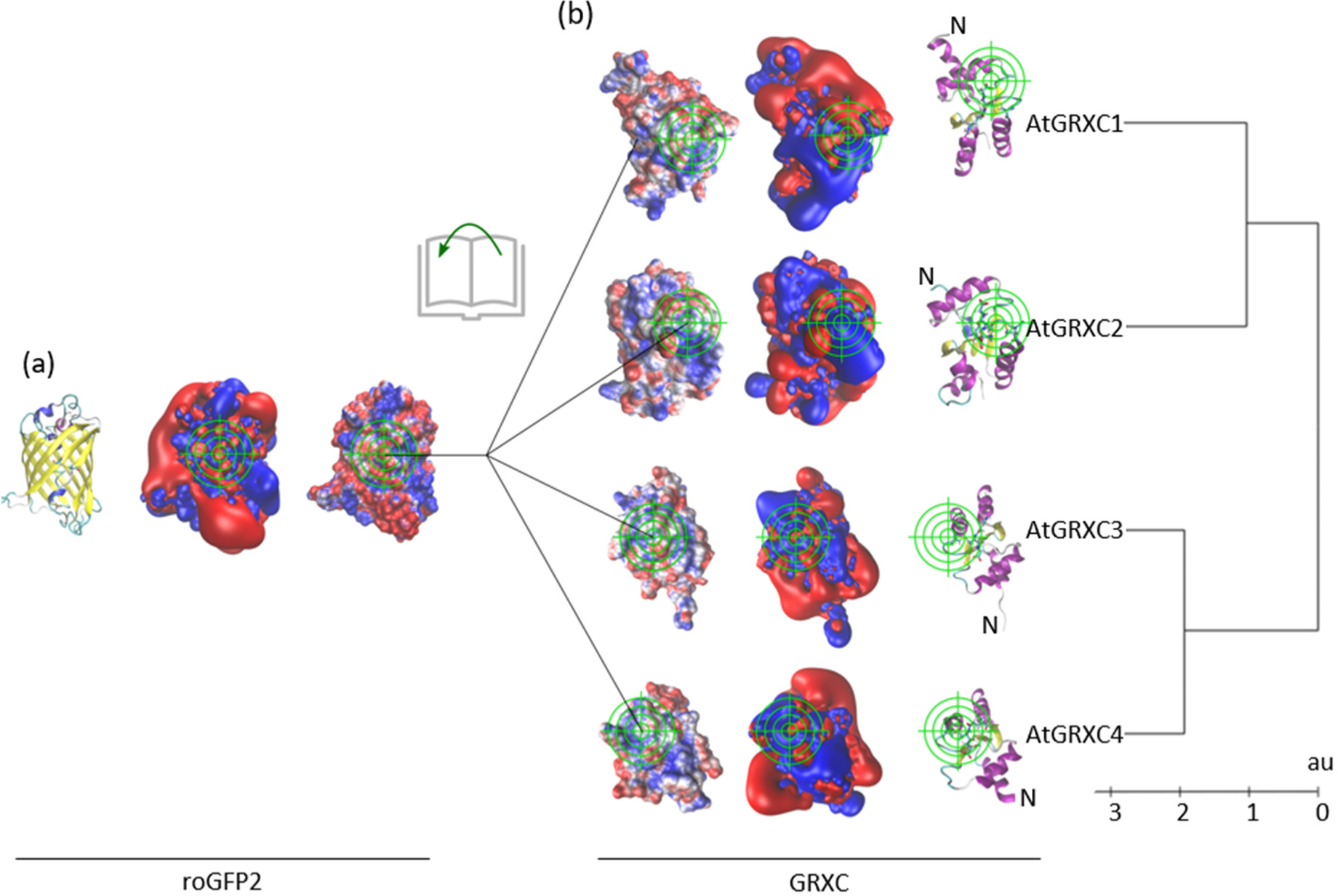
Clustering of class I glutaredoxins based on their electrostatic similarity and compatibility with roGFP2. The figure shows structural representations of roGFP2 (a) and GRXC1-4 (b) viewed from their interacting sites with reactive cysteines indicated by the green reticle. The proteins are depicted such that the reticle indicates Cys204 on roGFP2 and the N-terminal active site thiols for GRXCs with the axis oriented for a 180° attack characteristic for thiol-disulfide S_N_2 reactions. For the most compatible electrostatic interaction, GRXC3 and GRXC4 are rotated 180° compared to GRXC1 and GRXC2. For clarity, the N-termini of all GRX structures are indicated in the cartoon models (N). From left to right (or opposite for GRXCs) the proteins are depicted as cartoon models with α-helices colored in blue or purple and β-sheets in yellow, the electrostatic potential isosurfaces at ± 1 K T⋅e^-1^, and the electrostatic potential at ± 4 K T⋅e^-1^ mapped to the water-accessible surface of the proteins. Blue: positive, red negative potential. The pictures have been computed as outlined in the Methods section. The tree of electrostatic similarities of GRXs was calculated as described in Gellert et al. (2019).

## 4 | DISCUSSION

Class I GRXs with the canonical C(P/G/S)Y(C/S) active site are generally considered to mediate a number of different reactions including glutathionylation and deglutathionylation of substrate proteins, de-nitrosylation and de-persulfidation of protein thiols, dehydroascorbate reduction, and GSH-dependent peroxide reduction (Bedhomme et al., 2012; Begas et al., 2017; Moseler et al., 2021). While all these activities can be shown *in vitro*, the endogenous functions of class I GRXs are largely unknown and may vary on developmental stages, stress situation and subcellular localization of the respective GRXs. While localization of GRXC5 and its close paralog GRXS12 in plastids is well established (Couturier et al., 2011; Müller-Schüssele et al., 2021), localization of GRXC3 and GRXC4 is less clear. In both cases, bioinformatics algorithms only predict these proteins to likely enter the secretory pathway, but cannot make any more detailed predictions on their fate after transfer to the ER. Analysis of the primary sequence of GRXC1 shows a putative myristoylation site at the N-terminus, which may cause association of the protein with membranes. To narrow down the vast number of possible interactors of these proteins and to better understand their possible physiological function, we investigated the subcellular localization of all four non-plastidic class I GRXs and characterized their biochemical properties and potential functional redundancy of GRX isoforms that reside in the same compartment.

### 4.1 | GRXC1 is bound to membranes with an N-terminal myristoyl-group

Soluble proteins can be posttranslationally modified with fatty acids to alter their biochemical properties and especially the subcellular localization (Resh, 2016). N-terminal myristoylation is the common modification that directs proteins to membranes based on interaction of the lipophilic myristate residue with membrane lipids. Myristoylation requires a distinct motif MGxxxS/Txx that is present in Arabidopsis GRXC1 and many of its paralogous in other eudicot species. Indeed, GRXC1 was also found in the proteomic analysis of myristoylated proteins (Majeran et al., 2018). After initial co-translational cleavage of the N-terminal methionine, myristoyl-CoA:protein N-myristoyl transferase mediates the co-translational transfer of myristate from myristoyl-CoA to the N-terminal glycine via an amide bond (Breiman et al., 2016; Giglione et al., 2015). Myristoylation is necessary for membrane targeting but, due to the weak binding energy provided by the myristate moiety, insufficient for stable membrane anchoring (Majeran et al., 2018; Meinnel et al., 2020). Stable membrane anchoring, requires a second signal, which may be a polybasic cluster of amino acids for electrostatic interactions with phospholipids, hydrophobic amino acids along the membrane proximal protein surface, interaction with other proteins, or further modification with an additional palmitoyl residue (Running, 2014; Turnbull & Hemsley, 2017). Additional *S*-palmitoylation on a cysteine residue close to the myristoylation site also contributes to relocalization of the myristoylated protein to the plasma membrane (Traverso et al., 2013). All evidence gained so far with C-terminal roGFP2 fusions suggests localization of GRXC1 at endomembranes and the plasma membrane. While no additional cysteine close to the N-terminus is present, additional palmitoylation for plasma membrane targeting can be excluded. It is possible that strong overexpression of GRXC1-roGFP2 fusion proteins may lead to some artificial targeting to the plasma membrane. On the other hand, absence of residual soluble cytosolic protein indicates the high efficiency of the myristoylation site for recognition by the respective myristoylation machinery (Meinnel et al., 2020). While the exact positioning at distinct membranes or even membrane domains remains open for debate, all results strongly suggest that GRXC1 is always attached to the cytosolic leaflet of the membranes and thus displays its catalytic activity in the cytosol where it may potentially show some functional redundancy with GRXC2.

### 4.2 | GRXC3 and GRXC4 are type II transmembrane proteins in the secretory pathway

Expression of N- and C-terminal fusion proteins with roGFP2 consistently show that GRXC3 and GRXC4 are type II proteins in membranes along the early secretory pathway. This is consistent with earlier identification of both proteins in proteomes of the Golgi (Parsons et al., 2012) and GRXC4 in the ER (Nikolovski et al., 2012). It is noted, however, that both proteins have also been identified in the proteomes of several other locations along the secretory pathway and the apoplast (de Michele et al., 2016; Mitra et al., 2009; Nguyen-Kim et al., 2016; Takahashi et al., 2019; Zargar et al., 2015). Obviously, GFP fusion proteins may behave somewhat different from the native protein, which is exemplified by the fact, that the localization of GRXC3 and GRXC4 slightly shifted from the early ER for N-terminal fusions to the Golgi for C-terminal fusions. Such interference with correct protein positioning may happen in both directions with either an N-terminal GFP blocking transfer to the Golgi, or, alternatively, a C-terminal GFP disturbing retrieval from the Golgi to the ER. The same differential localization was observed earlier for glutathione peroxidase-like 3 (GPXL3), which is also a type II protein in the endomembrane system (Attacha et al., 2017). Similar to the observed relocalization to the Golgi, it can also not be excluded that the native protein at least in parts may be relocalized to the plasma membrane. Sorting proteins to their correct destinations are major functions of the secretory pathway in general, and the Golgi apparatus in particular (Lippincott-Schwartz & Phair, 2010). The rapid partitioning model assumes that different lipids undergo phase partitioning into two domains with different bilayer thickness (Bretscher & Munro, 1993). The predicted α-helical TMDs of 22 hydrophobic amino acids for GRXC3 and 21 amino acids for GRXC4 are about the average length that is sufficient to stretch a lipid bilayer (Cosson et al., 2013). It is noted though, that the second halves of the predicted TMDs of GRXC3 and GRXC4 contain several hydrophilic amino acids, which would necessarily reduce the hydrophobic interactions with adjacent lipids or may even lead to a shorter effective TMD. Mapping of the bilayer thickness in the ER membrane in yeast cells with fluorescent sensor proteins has shown discrete domains of increased bilayer thickness in the ER at ER–trans-Golgi contact sites. Thickening of the ER membrane contributes to the formation of lateral diffusion barriers, which restricted the passage of short, but not long, protein TMDs to budding cells (Prasad et al., 2020). Interestingly, GRXC4 has also been identified as a substrate of the vacuolar sorting receptor that in plant cells is thought to recognize proteins at the late Golgi or trans-Golgi network for vacuolar transport via the pre-vacuolar compartment (Shen et al., 2013). For such a transfer to work efficiently, however, the substrate protein GRXC4 would need to be soluble in the Golgi lumen rather than being membrane-anchored. Because the studied roGFP2 fusions of full-length proteins and of only the respective TMDs alone do not provide any hint that the TMD may get cleaved after recognition by the SEC61 translocon, we thus consider it more plausible that both GRXC3 and GRXC4 are anchored to the ER membrane with their N-termini to fulfill catalytic functions in the early secretory pathway.

### 4.3 | GRXs in the cytosol and in the secretory pathway show distinct biochemical properties

All four analyzed GRXs show a reductase activity in the classical HED assay, albeit with distinct difference between cytosolic and luminal isoforms. The higher turnover rates and catalytic efficiencies of GRXC3 and GRXC4 found here for Arabidopsis proteins is consistent with results for their paralogous from poplar (Couturier et al., 2013). This biochemical differentiation of cytosolic and luminal isoforms is also supported by a classification based on their electrostatic properties. While for several other representatives of the TRX superfamily analysis of the electrostatic properties have led to a revised functional classification that deviates from sequence-based approaches (Gellert et al., 2019), the classification identified here for GRX/roGFP2 interactions are fully consistent with the sequence-based phylogeny. Class I GRXs, however, are not only involved in mediating the reduction of substrates like tested in the HED assay but also the oxidation of suitable substrates, which are generally assumed to be protein thiols. A highly valuable experimental tool for further biochemical analysis of the respective activities is roGFP2, which can be reversibly reduced and oxidized by class I GRXs and can be exploited to study kinetic parameters of GRX activities (Begas et al., 2017; A. J. Meyer et al., 2007; Trnka et al., 2020). If electrostatic properties of the two interacting proteins would be critical, a similar differentiation between cytosolic and luminal GRXs would be expected. Surprisingly, however, the turnover rates for cytosolic and luminal GRXs were exactly reversed with GRXC3 and GRXC4 being more active in oxidation of roGFP2 than their cytosolic counterparts while GRXC1 and GRXC2 were more active in mediating the reduction reaction.

The differences in electrostatic properties may contribute to these differences in activities, as both cytosolic and luminal GRXs must adopt a different orientation towards the substrate (Figure 8) which must affect both affinity and reactivity towards the substrate. GRXC1 and GRXC2 display a higher affinity for S-glutathionylated substrates, *i.e.* the β-ME-S-SG formed in the HED assay, compared to GRXC3 and GRXC4. This was also reported for their counterparts from poplar The reduction of roGFP2 is supposedly initiated by a direct attack of GSH on the roGFP2 disulfide resulting on *S*-glutahionylated roGFP2. However, the equilibrium of this reaction must be on the site of the roGFP2 disulfide as no significant amounts of product are formed in the absence of GRXs, since roGFP2-SG has the same fluorescence properties as reduced roGFP2. The final reduction and the overall reaction must then by driven by the efficient reduction of the low amounts of roGFP2-SG by the GRXs. The oxidation is supposedly initiated by the formation of GRX-SG from reduced GRX and GSSG. Subsequently, GRX-SG glutathionylates roGFP2 to roGFP2-SG that, similar to the reduction reaction sequence, efficiently reacts to oxidized roGFP2. The reason for the opposite equilibria of reduction and oxidation reaction is the presence of 2 mM GSH in the reduction assay, while in the oxidation assay only minimal amounts (40 µM at maximum) of GSSG are present. The major discriminating factor for a preference for reduction and oxidation reactions may thus be the differences in affinity for the *S*-glutathionylated roGFP2 intermediate, especially if different orientations towards the substrate are required.

Sequence alignment for cytosolic and luminal GRXs highlights a number of distinct positional changes leading to a clear classification into four clades across multiple species. The most striking difference is an additional C-terminal cysteine that is present only in the subclades 1 and 2 representing the cytosolic isoforms. However, this cysteine is too far away from the active site and replacement by serine in *Pt*GRXC1 and *Pt*GRXC2 does not affect the reductase activity (Couturier et al., 2013). Several other residues that have changed between cytosolic and luminal isoforms are located in the α2-helix, which lies directly opposite of the active site motif on the α1-helix and includes several residues potentially involved in glutathione binding (Begas et al., 2017). It can be envisaged that these changes in total have subtle effects on the active site conformation and thus the catalytic activity. Further elucidation of these differences, however, requires the generation and detailed analysis of a large number of protein variants.

The distinct separation of luminal and cytosolic class I GRXs with two subclades in each group visible in trees generated on the basis of the surface electrostatic properties and the primary protein sequence likely resembles two genome duplication events with subsequent biochemical diversification. The very similar oxidoreductase activities in the two subclades suggests that the proteins in each compartment may potentially show functional redundancy. Indeed, some functional redundancy has been shown recently in *Saccharomyces cerevisiae* where a monocysteine variant of cytosolic *Sc*Grx2 was sufficient to rescue a lethal strain that is devoid of all endogenous GRXs and TRXs in the cytosol (Zimmermann et al., 2020). Furthermore, mutants of *Sc*GRX6 and *Sc*GRX7, which are both localized to the cis-Golgi, showed a more pronounced sensitivity to high temperature when both proteins are absent (Mesecke et al., 2008). While this raises further questions of why most class I GRXs have a CxxC active site motif, it indeed shows functional redundancy for some yet unknown essential physiological process.

Functional analysis of specific proteins would benefit from the availability of characteristic mutant phenotypes. Although null mutants for all four class I GRXs investigated here were isolated and confirmed, neither single nor *grxc1 grxc2* and *grxc3 grxc4* double mutants showed any distinct phenotype. For *grxc1 grxc2* this is particularly surprising because the respective double mutants had been described as non-viable (Riondet et al., 2012). Such lethality of double mutants would suggest at least partial redundancy of the two proteins, which is indeed the case from a biochemical viewpoint. However, the highly restricted expression pattern of *GRXC1* to later stages of embryogenesis and dry seeds (Figure S12), and in other tissues only in response to osmotic shock, together with the distinct association with membranes (Kilian et al., 2007; Sullivan et al., 2019; Waese et al., 2017), raises questions regarding functional redundancy. Functional diversification with GRXC1 being bound to membranes may reflect a co-evolutionary process during which specific substrate proteins at the membrane that are not accessible to the soluble GRXC2 with similar efficiency, have evolved. Possible substrate proteins of GRXC1 in the membrane could be TRXh7 and TRXh8, which are also myristoylated and reside predominantly on endomembranes where full-length TRX-GFP fusions accumulate on Golgi stacks (Traverso et al., 2013). On TRXh9, an additional cysteine close to the myristoylation site may get palmitoylated, which together with the myristoylation is assumed to mediate efficient and tight association with the plasma membrane (Traverso et al., 2013). Similar to the situation in the cytosol, duplication and subsequent diversification of ER-bound GRXs correlates with different expression patterns (Figure S12). Irrespective of the differentiation in expression patterns, the basic function of luminal GRXs may be glutathionylation or deglutathionylation during oxidative protein folding. The ability to monitor the luminal *E*_GSH_ with roGFP2-iL (Ugalde et al., 2022) is expected to provide further functional information on the luminal GRXs in future work.

## 5 | CONCLUSIONS

This study provides new insight in the localization of four class I GRXs in Arabidopsis and their biochemical properties. While GRXC1 and GRXC2 were known to be cytosolic already, new evidence provided here identifies GRXC1 as being anchored to membranes with an N-terminal myristoyl moiety. GRXC3 and GRXC4 are localized in the secretory pathway where they are both held in place by an N-terminal TMD. With the two remaining class I GRXs, GRXC5 and GRXS12, being established as plastidic proteins, this further confirms the absence of class I GRXs from mitochondria, which differs from the situation in yeast and mammalian cells. We show that all four studied GRXs are catalytically active as glutathione-dependent oxidoreductases that mediate reduction and oxidation of roGFP2 as a substrate protein. While the cytosolic GRXs are faster in catalyzing the reduction of roGFP2, both GRXs in the secretory pathway are faster in catalyzing the oxidation of roGFP2. The electrostatic properties of the four investigated GRXs match the sequence-based classification. With the opposing kinetics for reduction and oxidation of substrate proteins by cytosolic and luminal GRXs, it is, however, unlikely that differences in electrostatic GRX/roGFP2 interaction are accountable for the different kinetics. It is rather more likely that the differences are caused by subtle differences in overall protein structure that cause the equilibria of specific biochemical reaction steps to shift in one or the other direction. The differences result most likely from evolutionary adaptation to the different redox conditions in the two compartments with completely different redox conditions. In the absence of any distinct phenotypes of individual null mutants and also double-null mutants for the two class I GRXs and the fact that expansion of class I GRX is an evolutionary young process that occurred only after colonization of land, it might be possible that the function of these GRXs in the cytosol and the secretory pathway is more specific rather than very broad for regulation of a multitude of thiol proteins. While these functions need to be investigated in further work, the localization and biochemical characterization presented here should help to direct future functional studies.

## ACKNOWLEDGEMENTS

We thank Jean-Philippe Reichheld for providing the plasmids pET-16b GRXC1 and pET-16b GRXC2, and segregating *grxc1 grxc2* seeds, Aleksandra Wasilewska for providing a binary plasmid with GmMan1_1-49_-tdTomato and Jérémy Couturier for providing the plasmid pET-28a. Financial support by the Deutsche Forschungsgemeinschaft (DFG) (grant ME 1567/9-2 to A.J.M.) and the Alexander von Humboldt Foundation (A.M.) is gratefully acknowledged.

