## Supplemental Data for "Localization of four class I glutaredoxins in the cytosol and the secretory pathway and characterization of their biochemical diversification"

**Table S1** Primers used for genotyping, cloning, and determination of transcript abundance.

| **Name** | **Oligonucleotide sequence 5′ → 3′** |
|---|---|
| **Genotyping** | |
| LB-G (GABI-Kat) | ATATTGACCATCATACTCATTGC |
| LB-S (SALK) | ATTTTGCCGATTTCGGAAC |
| P1 | TTGCGGTCGAAACGTATATTC |
| P2 | ATGTACATCGCGCAGAAGAAG |
| P3 | ATCAAGCTCCACAACAAATGG |
| P4 | TCAGACCATTGGAAGAACCTG |
| P5 | ATCAAGCTCCACAACAAATGG |
| P6 | TCAGACCATTGGAAGAACCTG |
| P7 | AATTAAGGCCAAAGGAAAAGC |
| P8 | CCACCCACAACACTCTTTGAG |
| P9 | GAAGACTTGCGGTACTGTTCG |
| P10 | CAAATTGAACCCGCAAGATAG |
| P11 | TCGTCTCTGCTTATCCCGTT |
| P12 | TTGCCTTGCTTGTTGGTCTC |
| P13 | TCAGTTAACAGCGGAACCAAC |
| P14 | CGAATCTGTGTTTTTGGCTTC |
| **Determination of transcript abundance** | |
| P15 | ATGGGTTCAATGTTCAGTGGAAAC |
| P16 | TCTCCATCACTCTATCGCATCCAC |
| P17 | ATGGATTCGATTATGTGTGTTTCATC |
| P18 | AGTTTGATGTTGCATCACAGCCAC |
| P19 | ATGGTTGACCAGAGTCCTCG |
| P20 | AGCTCCAAGATCATCTGATCC |
| P21 | TCTCCGTCCTCTCTCTGATCAAG |
| P22 | ATGACAATGTTTAGATCTATCTCCA |
| P23 | ATCTACGGTATCATCTGATCCTC |
| P24 | TCCAACCATCTTCTCTTTCATCAAGC |
| **Gateway cloning** | |
| attB1 site | GGGGACAAGTTTGTACAAAAAAGCAGGCTTT |
| P25 | GGGGACAAGTTTGTACAAAAAAGCAGGCTTTATGGGTTCAATGTTCAGTGGAA |
| P26 | GGGGACAAGTTTGTACAAAAAAGCAGGCTTCGGTTCAATGTTCAGTGGAAACC |
| P27 | GGGGACAAGTTTGTACAAAAAAGCAGGCTTTATGGCTTCAATGTTCAGTGGAA |
| P28 | GGGGACCACTTTGTACAAGAAAGCTGGGTCTCAAAGTTGAGAAGAGTTATCTGCAA |
| P29 | GGGGACCACTTTGTACAAGAAAGCTGGGTCAAGTTGAGAAGAGTTATCTGCAATAG |
| P30 | GGGGACAAGTTTGTACAAAAAAGCAGGCTTTGCGATGCAGAAAGCTAAGGAGATCGAA |
| P31 | GGGGACAAGTTTGTACAAAAAAGCAGGCTTTATGGCGATGCAGAAAGCTAAGGAGATC |
| P32 | GGGGACCACTTTGTACAAGAAAGCTGGGTCTTAAGCAGAAGTTGTTGCAGTCTTT |
| P33 | GGGGACCACTTTGTACAAGAAAGCTGGGTCAGCAGAAGTTGTTGCAGTCTTTC |
| P34 | GGGGACAAGTTTGTACAAAAAAGCAGGCTTTGATTCGATTATGTGTGTTTCATCATG |
| P35 | GGGGACAAGTTTGTACAAAAAAGCAGGCTTTATGGATTCGATTATGTGTGTTTCATC |
| P36 | GGGGACAAGTTTGTACAAAAAAGCAGGCTTCTCTGGAATTTGCAGCAAGACT |
| P37 | GGGGACCACTTTGTACAAGAAAGCTGGGTCTTAAGCAGAAGTTGTTGCAGT |
| P38 | GGGGACCACTTTGTACAAGAAAGCTGGGTCTAAGCAAATTCCAGAAAAATTAC |
| P39 | GGGGACCACTTTGTACAAGAAAGCTGGGTCGCAAATTCCAGAAAAATTAC |

| | |
|---|---|
| P40 | GGGGACAAGTTTGTACAAAAAAGCAGGCTTTGCGAATTCTGTGTCAGCTTTC |
| P41 | GGGGACCACTTTGTACAAGAAAGCTGGGTCTCAACTTGCAGCAAGAAGCTTTT |
| P42 | GGGGACAAGTTTGTACAAAAAAGCAGGCTTTCATGGTTGACCAGAGTCCTCG |
| P43 | GGGGACCACTTTGTACAAGAAAGCTGGGTCACTTGCAGCAAGAAGCTTTTGCA |
| P44 | GGGGACCACTTTGTACAAGAAAGCTGGGTCTTAAGCTCCCGCAGAATTCGAAA |
| P45 | GGGGACCACTTTGTACAAGAAAGCTGGGTCAGCTCCCGCAGAATTCGAAA |
| P46 | GGGGACAAGTTTGTACAAAAAAGCAGGCTTTATGACAATGTTTAGATCTATCTCCAT |
| P47 | GGGGACAAGTTTGTACAAAAAAGCAGGCTTTGCTGCTTCGTCCCCAGAA |
| P48 | GGGGACCACTTTGTACAAGAAAGCTGGGTCCTAGAGTTCAGCTTCTTTGTTCC |
| P49 | GGGGACCACTTTGTACAAGAAAGCTGGGTCGAGTTCAGCTTCTTTGTTCCC |
| P50 | GGGGACCACTTTGTACAAGAAAGCTGGGTCTTAAGAAGAAACCATAGAAATGA |
| P51 | GGGGACCACTTTGTACAAGAAAGCTGGGTCAGAAGAAACCATAGAAATGA |
| **Restriction enzyme-based cloning** | |
| P52 | CCCCCCATGGCGTCCCCAGAAGCCGACTTT |
| P53 | CCCCTCGAGGAGTTCAGCTTCTTTGTTCCCGGA |

**Table S2** Primer pairs used for amplification of full length coding sequence (CDS), CDS without transmembrane domains (TMDs), and TMDs alone.

| Purpose | Gene | Primer | | Entry Plasmid |
|---|---|---|---|---|
| | | **Forward** | **Reverse** | |
| Full length CDS **with stop** codon for fusion of roGFP2 to the **N-termini** | GRXC1 | P26 | P28 | pDONR201 |
| | GRXC2.1 | P30 | P32 | |
| | GRXC2.2 | P34 | P32 | |
| | GRXC3 | P40 | P41 | |
| | GRXC4 | P46 | P48 | |
| Full length **without** stop codon for fusion of roGFP2 to the **C-termini** | GRXC1 | P25 | P29 | pDONR207 |
| | GRXC1 (G2A) | P27 | P29 | |
| | GRXC2.1 | P31 | P33 | |
| | GRXC2.2 | P35 | P33 | |
| | GRXC3 | P40 | P43 | |
| | GRXC4 | P46 | P49 | |
| CDS without TMD **with stop** codon for fusion of roGFP2 to the **N-termini** | GRXC3 | P40 | P41 | pDONR201 |
| | GRXC4 | P47 | P48 | |
| CDS without TMD and **without stop** codon for fusion of roGFP2 to the **C-termini** | GRXC3 | P40 | P42 | pDONR207 |
| | GRXC4 | P47 | P49 | |
| TMD **with stop** codon for fusion of roGFP2 to the **N-termini** | GRXC2.2 | P35 | P38 | pDONR201 |
| | GRXC3 | P42 | P44 | |
| | GRXC4 | P46 | P50 | |
| TMD **without stop** codon for fusion of roGFP2 to the **C-termini** | GRXC2.2 | P35 | P39 | pDONR207 |
| | GRXC3 | P42 | P45 | |
| | GRXC4 | P46 | P51 | |

**Table S3** Gene identifiers

| Code used in this publication | Gene identifier |
|---|---|
| *At*GRXC1 | AT5G63030 |
| *Pt*GRXC1, 2 | Potri.012G082800 |
| *Pt*GRXC1, 1 | Potri.015G078900 |
| *Mt*GRXC1 | Medtr3g077560 |
| *Sl*GRXC1 | Solyc03g112770 |
| *St*GRXC1 | Soltu.DM.03G026520 |
| *Vv*GRXC1 | VIT_217s0000g08670 |
| *At*GRXC2 | AT5G40370 |
| *Pt*GRXC2 | Potri.001G347700 |
| *OsGRXC2, 1* | LOC_Os04g42930 |
| *Mt*GRXC2 | Medtr7g035245 |
| *Sl*GRXC2 | Solyc06g005260 |
| *St*GRXC2 | Soltu.DM.06G004540 |
| *Vv*GRXC2 | VIT_214s0066g00960 |
| *OsGRXC2, 2* | LOC_Os02g40500 |
| *Hv*GRXC2 | HORVU.MOREX.r3.2HG0180500 |
| *At*GRXC3 | AT1G77370 |
| *Pt*GRXC3 | Potri.007G017300 |
| *Os*GRXC3 | LOC_Os02g43180 |
| *Mt*GRXC3 | Medtr5g021090 |
| *Sl*GRXC3 | Solyc02g084660 |
| *St*GRXC3 | Soltu.DM.02G024470 |
| *Vv*GRXC3 | VIT_204s0023g02800 |
| *Hv*GRXC3 | HORVU.MOREX.r3.6HG0599220 |
| *At*GRXC4 | AT5G20500 |
| *Pt*GRXC4 | Potri.018G133400 |
| *Os*GRXC4 | LOC_Os06g44910 |
| *Mt*GRXC4 | Medtr2g038560 |
| *Sl*GRXC4 | Solyc11g069860 |
| *St*GRXC4 | Soltu.DM.11G023100 |
| *Vv*GRXC4 | VIT_211s0052g00500 |
| *Hv*GRXC4 | HORVU.MOREX.r3.7HG0740050 |
| *At*GRXC5 | AT4G28730 |
| *Mt*GRXC5 | Medtr1g069255 |
| *At*GRXS12 | AT2G20270 |
| *Os*GRXS12 | LOC_Os08g45140 |
| *Hv*GRXS12 | HORVU.MOREX.r3.7HG0693390 |
| *Pt*GRXS12 | Potri.002G254100 |
| *Sl*GRXS12 | Solyc06g083690 |
| *St*GRXS12 | Soltu.DM.06G034090 |
| *Vv*GRXS12 | VIT_207s0005g04860 |

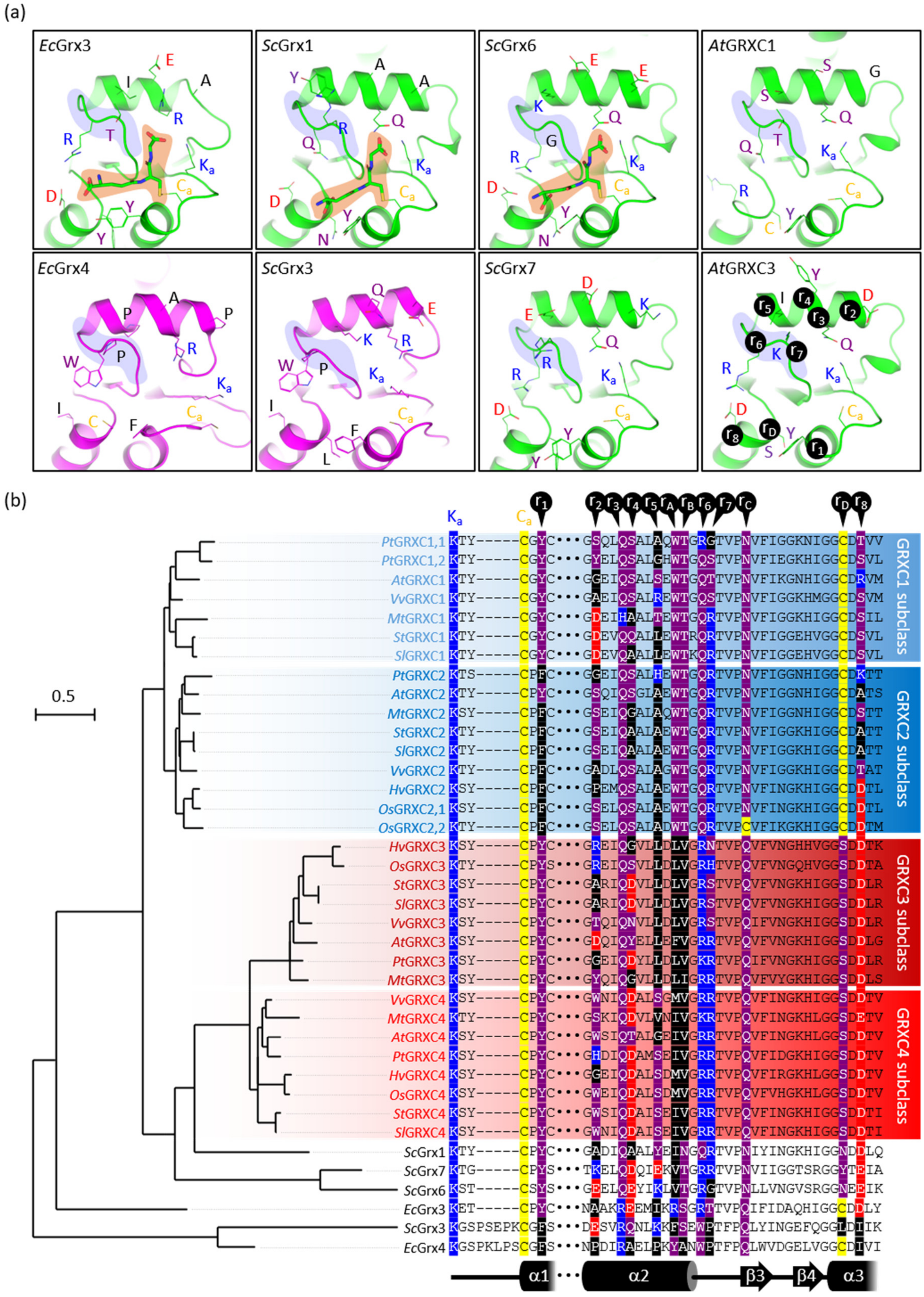


**FIGURE S1** Structurally and sequentially conserved features of representative glutaredoxins from *E. coli*, yeast and plants. (a) Superimposition of representative glutaredoxin structures. Homology models of Arabidopsis GRXC1 and GRXC3 were obtained through the Phyre2 webserver (Kelley et al., 2015) with default settings and the GRXC1 and GRXC3 amino acid sequences as input. A structural model of *Saccharomyces cerevisiae* (*Sc*) Grx7 was obtained from Liedgens et al. (2020). Crystal structures of *Escherichia coli* (*Ec*) and yeast (*Sc*) class I (green) and class II (magenta) glutaredoxins were obtained from the Protein Data Bank (PDB) using the following PDB codes. *Ec*Grx3: 3grx; *Ec*Grx4: 1yka; *Sc*Grx1: 3c1s; *Sc*Grx3: 5y4u; *Sc*Grx6: 3l4n. The conserved active-site cysteine (Ca) and lysine (Ka) residue, potential glutathione-interacting residues r1–

8, as defined in Liedgens et al. (2020) and a conserved S/C position (rD) at the glutaredoxin C-terminus are shown as sticks on the cartoon models. A co-crystallized glutathione (GSH) molecule in *Ec*Grx3, *Sc*Grx1, *Sc*Grx6 is highlighted in orange. The 'basic loop' region following helix α2 is highlighted in blue. (b) Sequence alignment and phylogeny of glutaredoxins shown in Figure 1 and in panel (a). The scale bar represents 0.5 substitutions per amino acid position. Positions r1–8, as defined in Liedgens et al. (2020), the conserved active-site cysteine (Ca) and lysine (Ka) residue and positions rA-D are color-coded based on the amino acid properties. Positions rA-D are largely conserved within GRXC1/C2 and GRXC3/C4 subclasses, respectively, but different between the GRXC1/C2 (light and dark blue) and GRXC3/C4 (dark and light red) subclasses.

GOUY M., GUINDON S. & GASCUEL O. *Molecular Biology Evol*. 27:221–224. (2010)

KELLEY, L., MEZULIS, S., YATES, C.*et al. Nature Protocos* **10**, 845–858 (2015).

LIEDGENS L, ZIMMERMANN J, WÄSCHENBACH L, GEISSEL F, LAPORTE H, GOHLKE H, MORGAN B, DEPONTE M. *Nature Communnication*. 11:1725. (2020)

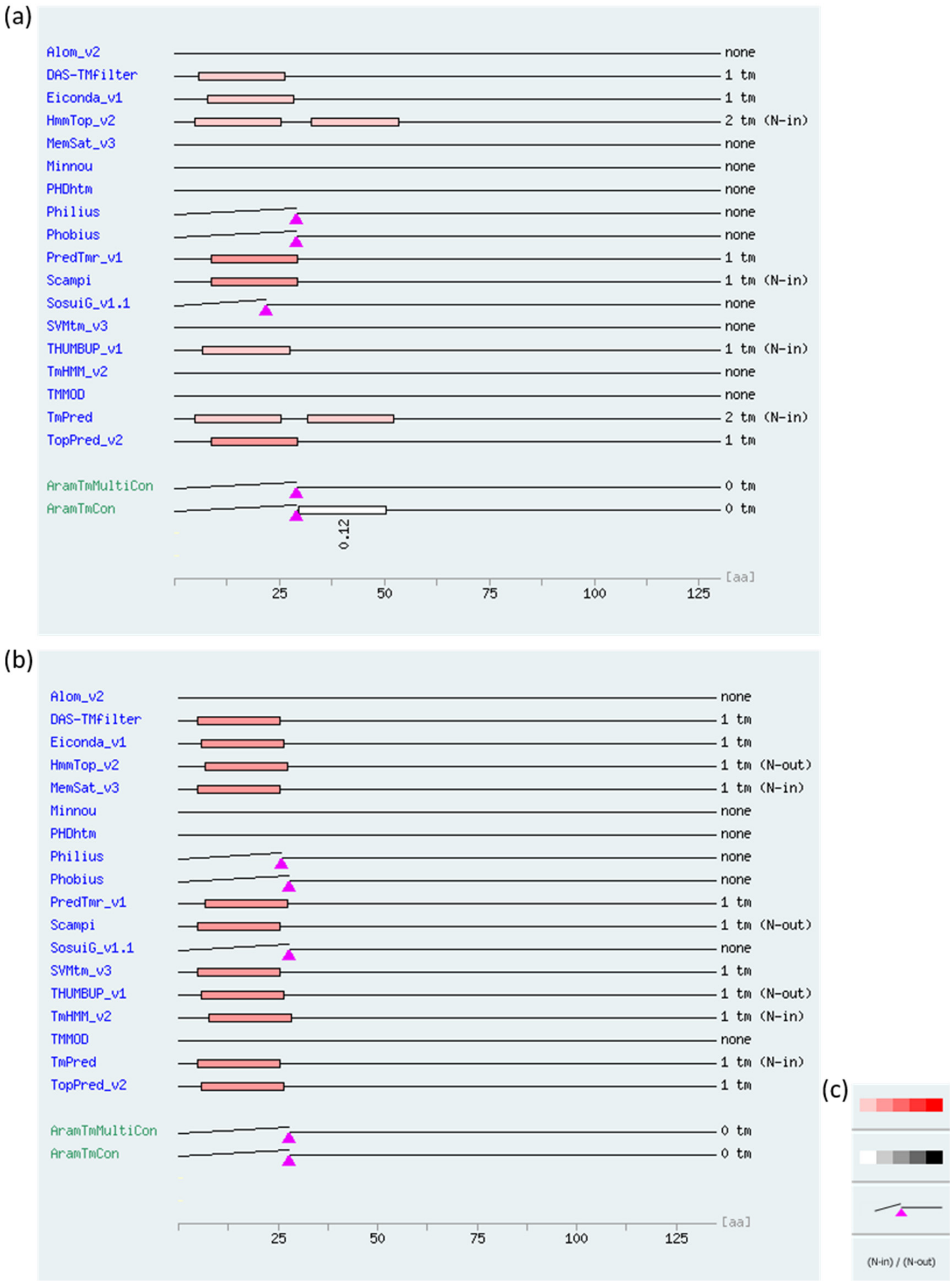


**Figure S2** Graphical representation of predicted topology models for GRXC3 and GRXC4 from *Arabidopsis thaliana*. (a,b) Visualization of putative transmembrane domains (TMDs) and cleavable target peptides for GRXC3 (a) and GRXC4 (b) by different bioinformatics algorithms. The figures show results generated by ARAMEMNON (http://aramemnon.uni-koeln.de/) (Schwacke *et al.*, 2003; Schwacke & Flügge, 2018) and include two consensus predictions AramTmCon (based on individual predictions for the protein) and AramTMMultiCon, that also takes into account predictions for orthologous proteins. Hydrophobic domains interpreted as $\alpha$-helical TMDs are shown as boxes with color saturation indicating the degree of hydrophobicity. Triangles indicate cleavage sites for a predicted target peptide. *N-in* indicates the N-termini facing the cytosol while *N-out* indicates extracytosolic orientation.

SCHWACKE R, SCHNEIDER A, VAN DER GRAAFF E, FISCHER K, CATONI E, DESIMONE M, FROMMER WB, FLÜGGE UI, KUNZE R. 2003.Plant Physiol, Jan;131(1):16-26.

SCHWACKE R, FLÜGGE UI. 2018. Methods Molecular Biology, 1696:249-259.

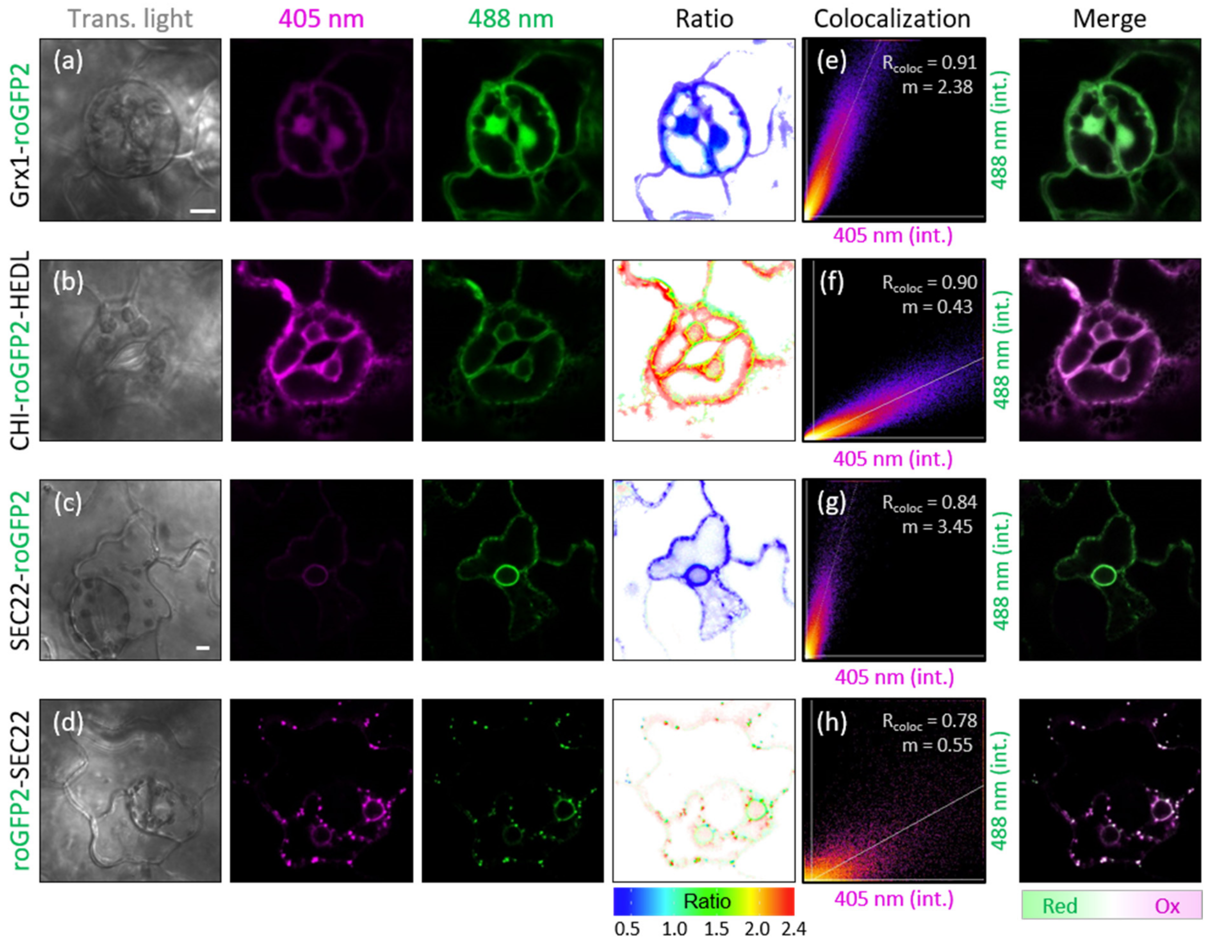


**Figure S3** Redox state of cytosolic and ER-targeted roGFP2 for *in vivo* analysis of membrane protein orientation. (a,b) Confocal microscopy images of guard cells from 7-day-old Arabidopsis seedlings stably expressing Grx1-roGFP2 (Marty *et al.*, 2009) in the cytosol (a) or $CHI_{TP}$-roGFP2-HDEL (Schwarzländer *et al.*, 2008) targeted to the ER-lumen (b). (c,d) Leaf epidermal cells from tobacco (*Nicotiana tabacum*) transiently expressing roGFP2 fused to the N-terminus (c) or C-terminus (d) of the ER-targeted SEC22 (Brach *et al.*, 2009). Images in the second and third column show roGFP2 fluorescence collected at 505–530 nm after excitation at either 405 nm (magenta) or 488 nm (green). The level of sensor oxidation is also displayed as the 405 nm/488 nm ratio, false-colored from blue to red for reduced or oxidized sensor, respectively. Bar = 5 µm. Note that the localization of N- and C-terminally tagged SEC22 differs with roGFP2-SEC22 localizing preferentially in Golgi stacks. Also, the roGFP2 in this case appears not always fully oxidized, probably due to formation of homodimers with two intermolecular disulfide that lead to a pseudo-reduced state of the chromophore (Sarkar *et al.*, 2013). The scatter plots show fluorescence intensities for excitation at 405 nm (x-axis) and 488 nm (y-axis), respectively, plotted against each other for each pixel. Fluorescence intensities of both channels were evaluated according to their Pearson correlation coefficient above the threshold ($R_{coloc}$). The gradient (m) clearly separates reduced (m > 1) Grx1-roGFP2 and roGFP2-SEC22 from oxidized (m < 1) $CHI_{TP}$-roGFP2-HDEL and SEC22-roGFP2. Based on this binary readout the obtained color after merging the two channels also reliably indicates the oxidation level of roGFP2 with green for reduced and magenta for oxidized roGFP2, respectively.


MARTY, L., SIALA, W., SCHWARZLÄNDER, M., FRICKER, M., WIRTZ, M., SWEETLOVE, L., MEYER, Y., MEYER, A., REICHHELD, J., & HELL, R. 2009. Proceedings of the National Academy of Sciences of the United States of America, 106(22), 9109–9114.BRACH, T., SOYK, S., MÜLLER, C., HINZ, G., HELL, R., BRANDIZZI, F. & MEYER, A. J. 2009. The Plant Journal, 57, 534-541.

SARKAR, D., EDWARDS, S., MAUSER, J., SUAREZ, A., SEROWOKY, M., HUDOK, N., HUDOK, P., NUÑEZ, M., WEBER, C., LYNCH, R., MIYASHITA, O. AND TSAO, T. 2013. Biochemistry, 52 (19), 3332-3345

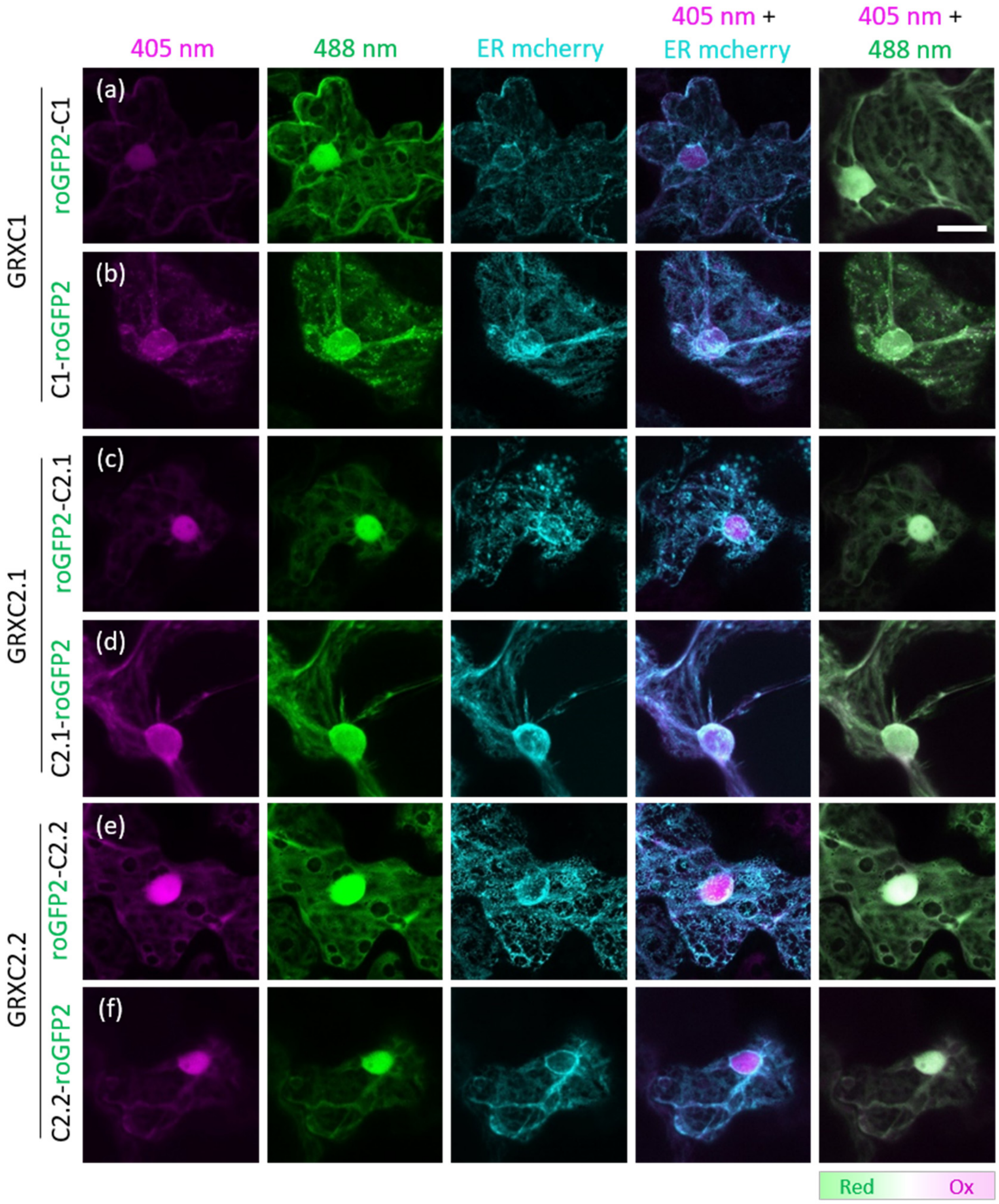


**Figure S4** Subcellular localization of GRXC1, GRXC2.1 and GRXC2.2 in tobacco. (a-f) Representative confocal microscopy images of pavement cells in the leaf epidermis transiently expressing roGFP2 fusions to the N- and C-termini of GRXC1 (a,b), GRXC2.1 (c,d) and GRXC2.2 (e,f). Images show roGFP2 fluorescence collected at 505–530 nm after excitation with either 405 nm (magenta) or 488 nm (green). For ER colocalization, leaves were co-infiltrated with the marker AtWAK2$_{TP}$-mCherry-HDEL. The green color obtained after merging the 405 nm and 488 nm roGFP2 channels indicates roGFP2 in all constructs as reduced. Bar = 20 µm.

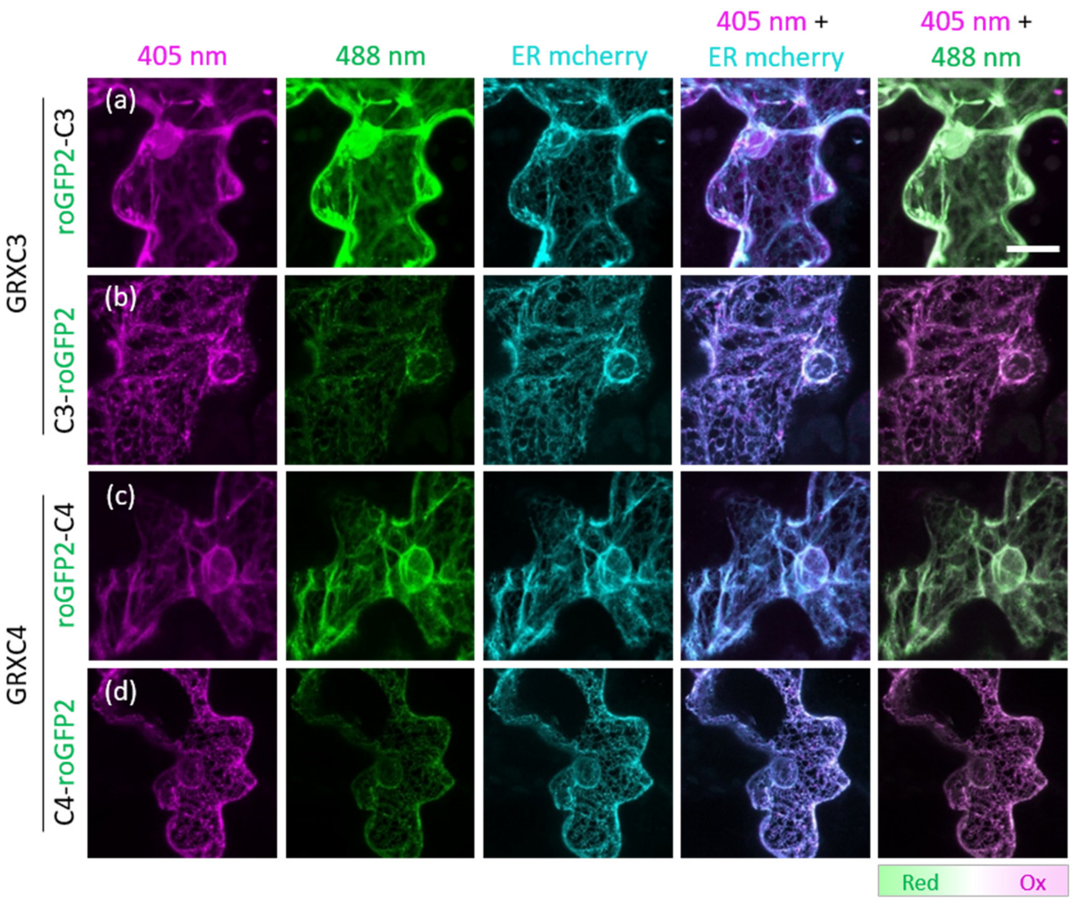


**Figure S5** Subcellular localization of GRXC3 and GRXC4 in tobacco. (a-d) Representative confocal microscopy images of pavement cells in the leaf epidermis transiently expressing roGFP2 fusions to the N- and C-termini of GRXC3 (a,b) or GRXC4 (c,d). Images show roGFP2 fluorescence collected at 505–530 nm after excitation with either 405 nm (magenta) or 488 nm (green). To test for localization in the ER, the respective roGFP2 fusion constructs were co-expressed with the ER marker AtWAK2$_{TP}$-mCherry-HDEL. The colors obtained after merging the 405 nm and 488 nm channels indicate the oxidation level of roGFP2 on a scale from green to magenta for reduced to oxidized, respectively. Bar = 20 µm.

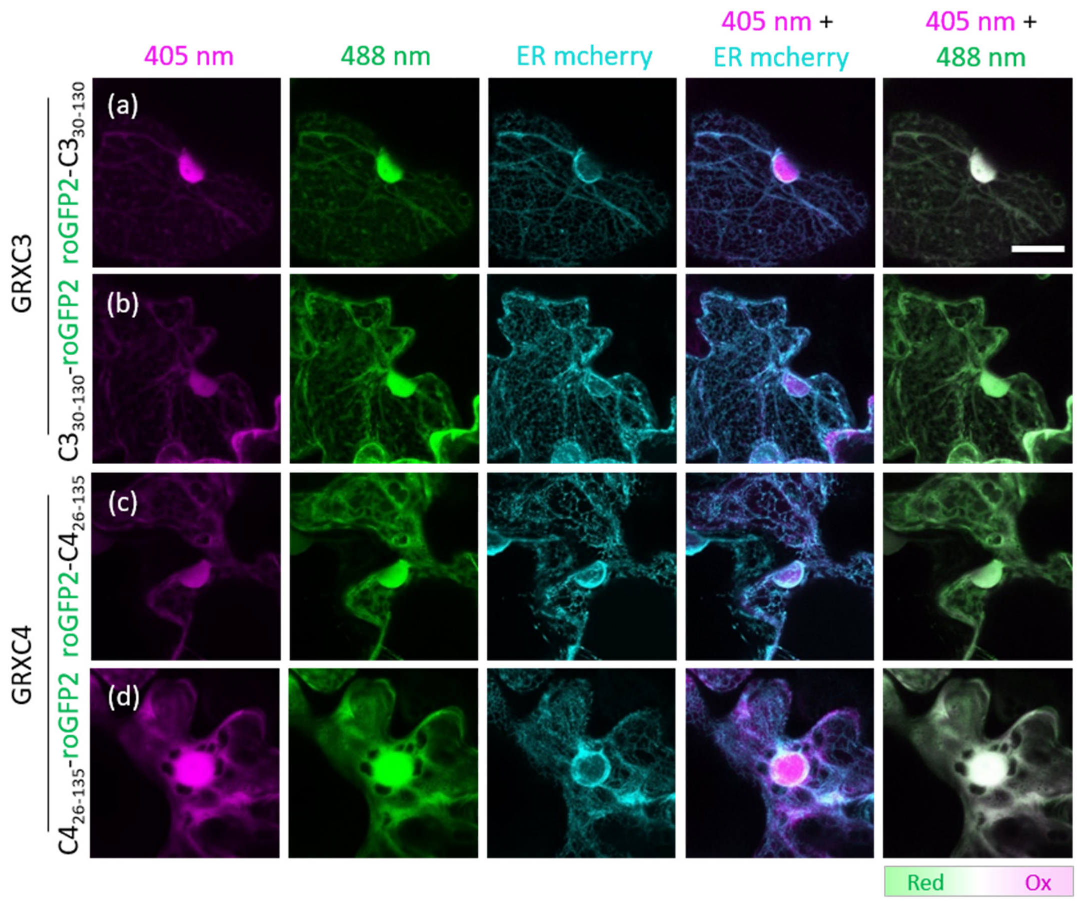


**Figure S6** Subcellular localization of glutaredoxins GRXC3$_{30\text{-}130}$ and GRXC4$_{26\text{-}135}$ lacking their N-terminal transmembrane domains in tobacco. (a-d) Representative confocal microscopy images of pavement cells in the leaf epidermis transiently expressing roGFP2 fusions to the N- and C-termini of GRXC3$_{\mathbf{30\text{-}130}}$ (a,b) or GRXC4$_{\mathbf{26\text{-}135}}$ (c,d). Images show roGFP2 fluorescence collected at 505–530 nm after excitation with either 405 nm (magenta) or 488 nm (green). For counterstaining of the ER, leaves were co-infiltrated with the marker AtWAK2$_{TP}$-mCherry-HDEL. The colors obtained after merging the 405 nm and 488 nm roGFP2 channels indicate the oxidation level of roGFP2 on a scale from green to magenta for reduced to oxidized, respectively. Bar = 20 µm.

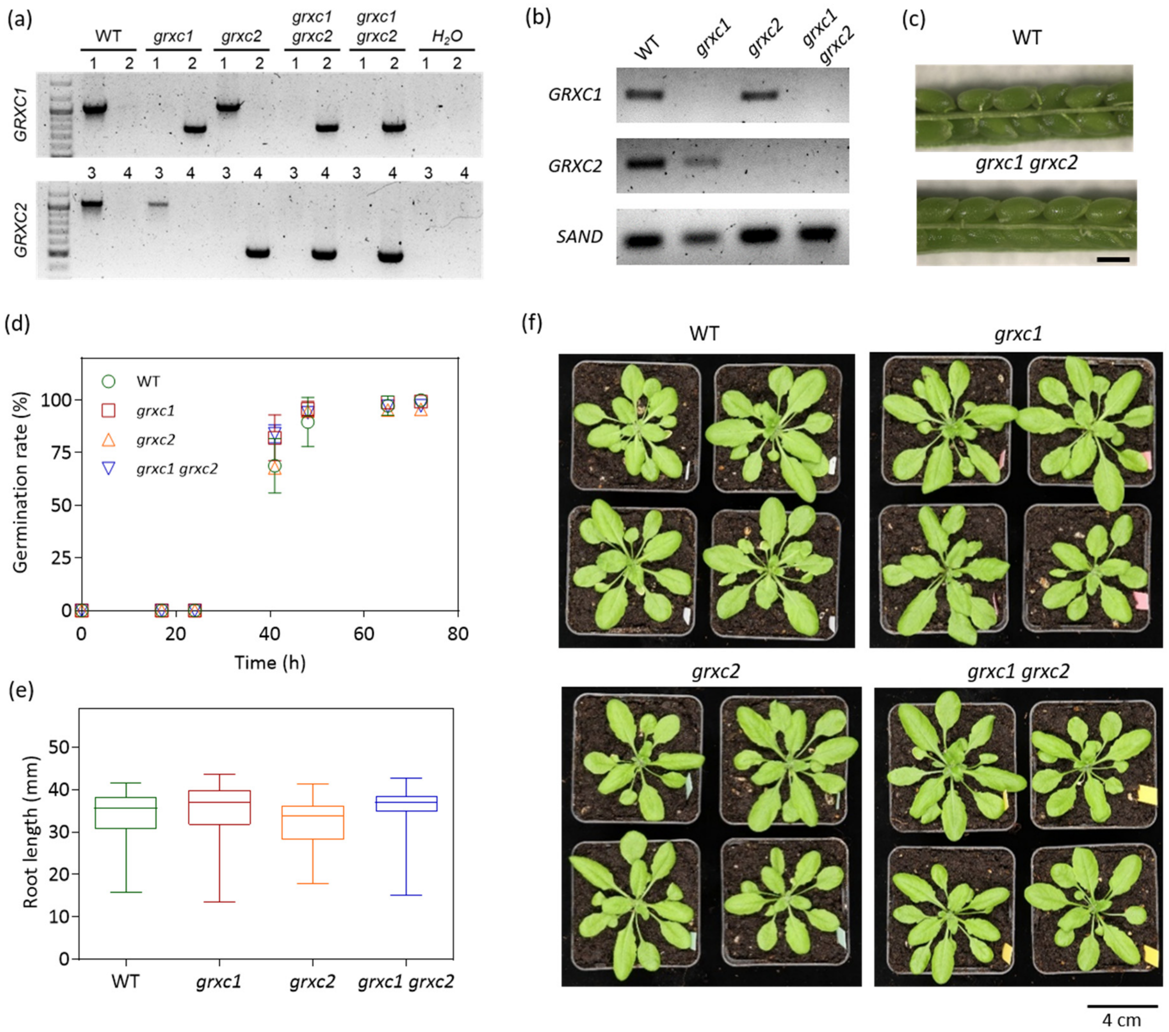


**Figure S7** Isolation and phenotypic characterization of a *grxc1 grxc2* double mutant. (a) Genotyping for *grxc1* (upper panel) using the primer combinations P11 + P12 (1) and P12 + LB-G (2) and for *grxc2* (lower panel) with the primer combinations P13 + P14 (3) and P14 + LB-S (4) as indicated above the respective lanes. Wild type and homozygous single mutants were used as controls. (b) Assessment of expression levels for *GRXC1* and *GRXC2*. Total RNA extracted from 2-week-old seedlings was used for RT-PCR with a forward primer annealing to the start of the coding region and an exon-exon spanning reverse primer annealing to parts of exon three and exon four in the coding region of *GRXC1* (P15 + P16) or to parts of exon two and exon three in the coding region of *GRXC2* (P17 + P18). (c) Opened siliques from wild-type and double homozygous *grxc1 grxc2* plants. Bar = 500 µm. (d) Germination rate of WT, *grxc1* and *grxc2* single mutants, and the respective *grxc1 grxc2* double mutant. All seeds were initially stratified at 4 °C in the dark for 1 day (*n* = 3-4 with 36–115 seeds each; means ± SD). Germination was assessed on the basis of radicle emergence. (e) Primary root length of 8-day-old seedlings grown on vertical plates under long-day conditions. (*n* = 33-60; the box plot shows the median as center line with the box for the first to the third quartile and whiskers indicating min and max values). (f) Phenotypic comparison of 4-week-old plants grown on soil under long-day conditions.

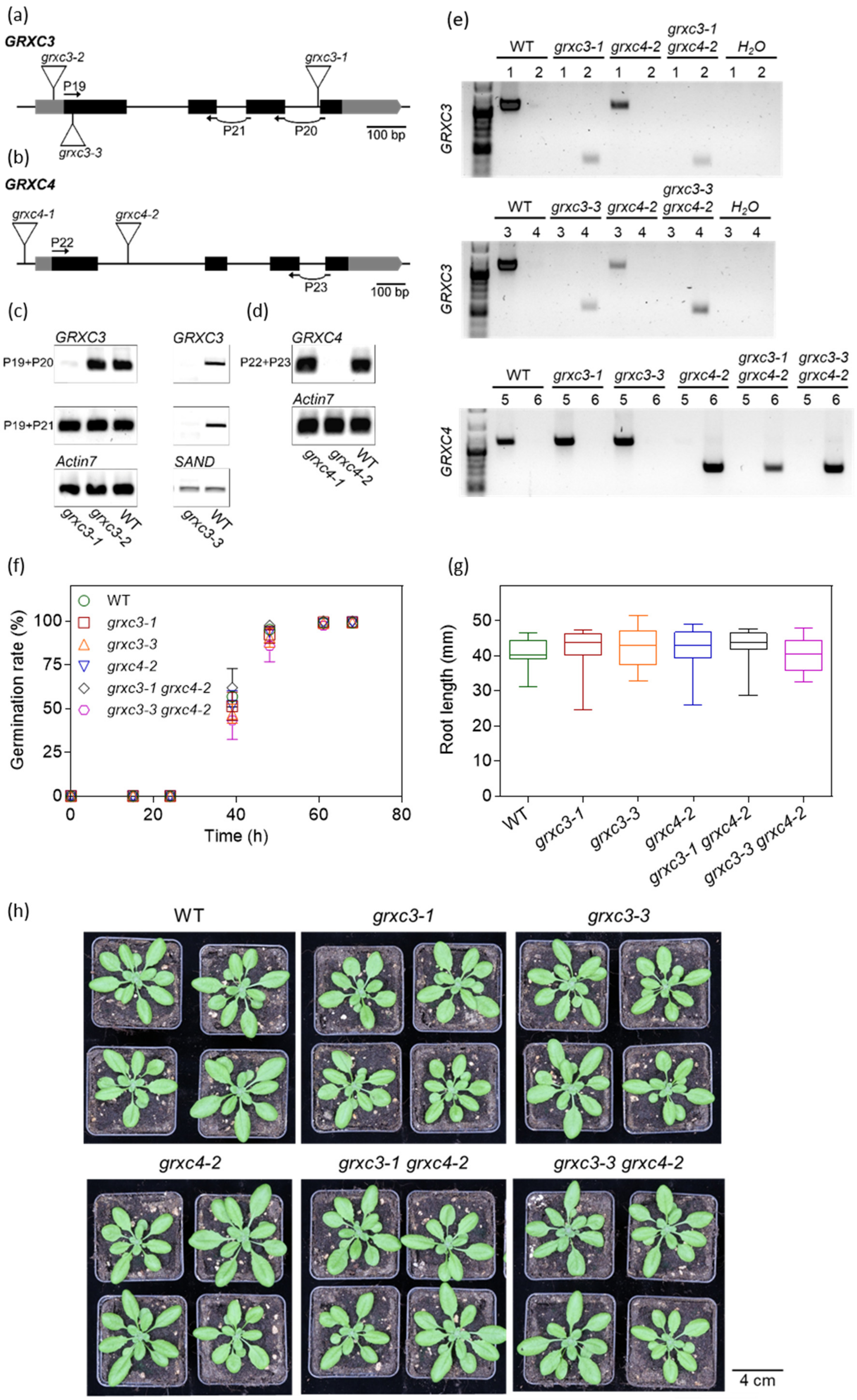
(a)
GRXC3
grxc3-2
P19
grxc3-1
P21
P20
100 bp
grxc3-3
(b)
GRXC4
grxc4-1
grxc4-2
P22
P23
100 bp
(c)
GRXC3
P19+P20
P19+P21
Actin7
grxc3-1
grxc3-2
WT
GRXC3
SAND
grxc3-3
WT
(d)
GRXC4
P22+P23
Actin7
grxc4-1
grxc4-2
WT
(e)
WT
grxc3-1
grxc4-2
grxc3-1 grxc4-2
$H_2O$
1 2
GRXC3
grxc3-3
grxc3-3 grxc4-2
3 4
GRXC3
5 6
GRXC4
(f)
Germination rate (%)
Time (h)
WT
grxc3-1
grxc3-3
grxc4-2
grxc3-1 grxc4-2
grxc3-3 grxc4-2
(g)
Root length (mm)
WT
grxc3-1
grxc3-3
grxc4-2
grxc3-1 grxc4-2
grxc3-3 grxc4-2
(h)
WT
grxc3-1
grxc3-3
grxc4-2
grxc3-1 grxc4-2
grxc3-3 grxc4-2
4 cm

**Figure S8** Isolation and phenotypic characterization of *grxc3* and *grxc4* mutants*.* (a,b) Physical maps of *GRXC3* (At1g77370) and *GRXC4* (At5g20500). Introns are represented as lines and exons as boxes. Both UTRs are depicted in gray. The primers used for RT-PCR are indicated by numbered arrows. The T-DNA insertions of the mutant lines are shown as inverted triangles. (c,d) Analysis of *GRXC3 (C)* and *GRXC4* (D) expression. For RT-PCR, a forward primer annealing to the start of the coding region and exon-exon spanning reverse primer annealing to exon two and exon three in the coding region of *GRXC3* (P19 + P20) or to exon three and exon four in the coding region of *GRXC3 (P19 + P21) or GRXC4* (P22 + P23) were used. (e) Representative gel showing the genotyping for GRXC3 (upper panel) using the primer combination P1 + P2 (1) and P1 + LB-G (2) for *grxc3-1* or P5 + P6 (3) and P5 + LB-S (4) for *grxc3-3* and genotyping for GRXC4 (lower panel) using the primer combination P9 + P10 (5) and P9 + LB-S (6). Wild type and single mutants were used as control. (f) Germination rate of WT, *grxc3, grxc4* single mutants and the respective double mutants. All seeds were initially stratified at 4 °C in the dark for 1 day (*n* = 3 with 76–118 seeds each; means ± SD). Germination was assessed with the emergence of the radicle. (g) Primary root length of 8-day-old mutants compared to WT on vertical plates under long-day conditions. (*n* = 19-21; the box plot shows the median as center line with the box for the first to the third quartile and whiskers indicating min and max values). (h) Phenotypic comparison of 4-week-old plants grown on soil under long-day conditions.

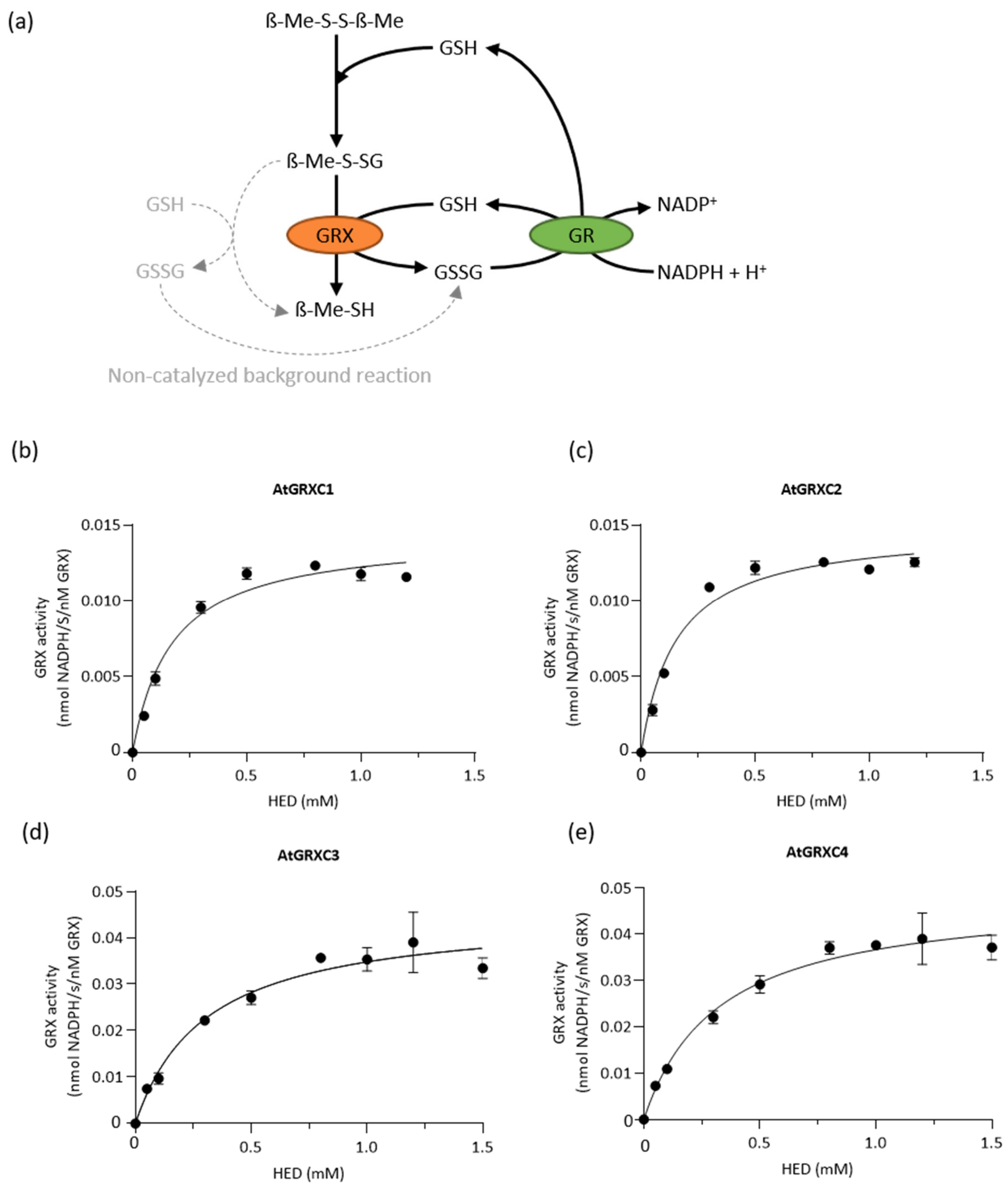


**Figure S9** HED assay for cytosolic and luminal class I glutaredoxins. (a) Scheme of the sequential reactions taking place. GRX activity was followed as the oxidation of NADPH (decrease in $Abs_{340}$) in a coupled reaction with GSH and glutathione disulfide reductase (GR), where GRX catalyzes the reduction of ß-ME-S-SG by GSH. (b-e) Michaelis-Menten enzyme kinetics for 40 nM GRXC1 (b), 40 nM GRXC2 (c), 10 nM GRXC3 (d) and 10 nM GRXC4 (e). Data shown are mean ± SD from a single experiment with three technical replicates from one protein purification and dilution *n* = 3.

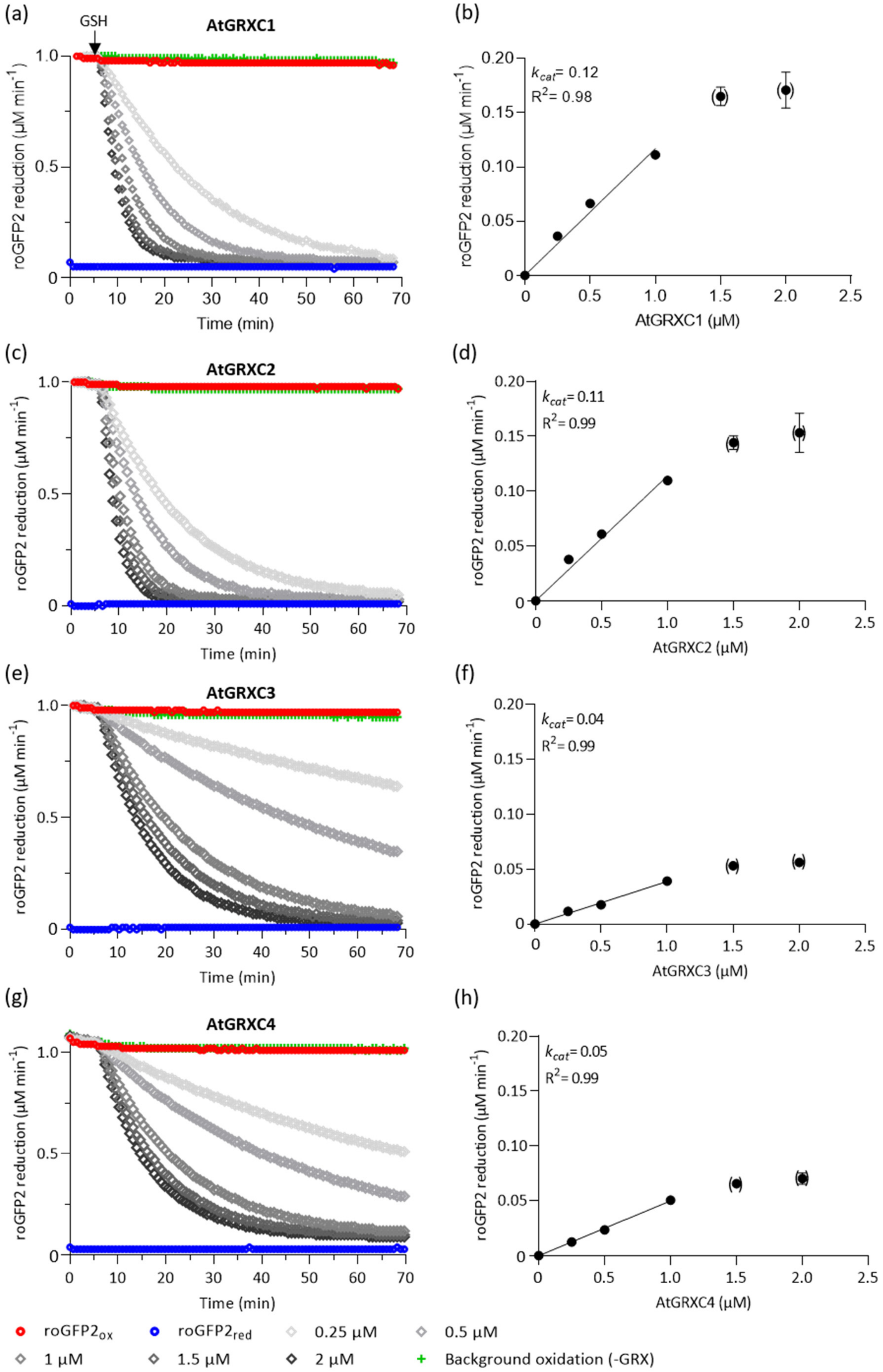
(a)
GSH
AtGRXC1
roGFP2 reduction (μM min-1)
Time (min)
(b)
kcat= 0.12
R2= 0.98
roGFP2 reduction (μM min-1)
AtGRXC1 (μM)
(c)
AtGRXC2
roGFP2 reduction (μM min-1)
Time (min)
(d)
kcat= 0.11
R2= 0.99
roGFP2 reduction (μM min-1)
AtGRXC2 (μM)
(e)
AtGRXC3
roGFP2 reduction (μM min-1)
Time (min)
(f)
kcat= 0.04
R2= 0.99
roGFP2 reduction (μM min-1)
AtGRXC3 (μM)
(g)
AtGRXC4
roGFP2 reduction (μM min-1)
Time (min)
(h)
kcat= 0.05
R2= 0.99
roGFP2 reduction (μM min-1)
AtGRXC4 (μM)
roGFP2ox
roGFP2red
0.25 μM
0.5 μM
1 μM
1.5 μM
2 μM
Background oxidation (-GRX)

**Figure S10** Activity of class I GRXs in the reduction of roGFP2. (a, c, e, g) GSH dependent reduction of 1 µM oxidized roGFP2. All GRXs were tested from 0.25 µM (light grey) to 2 µM (dark grey). The background reaction recorded in the absence of GRX is depicted in green. Full reduction or oxidation were determined by incubating roGFP2 with 10 mM $DTT_{red}$ or 10 mM $DTT_{ox}$, respectively. The arrows mark the time point at which the assays were started by addition of 2 mM GSH. (b, d, f, h) Concentrations of the GRXs used versus activity. All curves were fitted with a linear regression for data up to 1 µM GRX to determine $k_{cat}$ values. The data points in parenthesis were excluded from the analyses. Data shown are mean ± SD from a single experiment with three technical replicates from one protein purification and dilution.

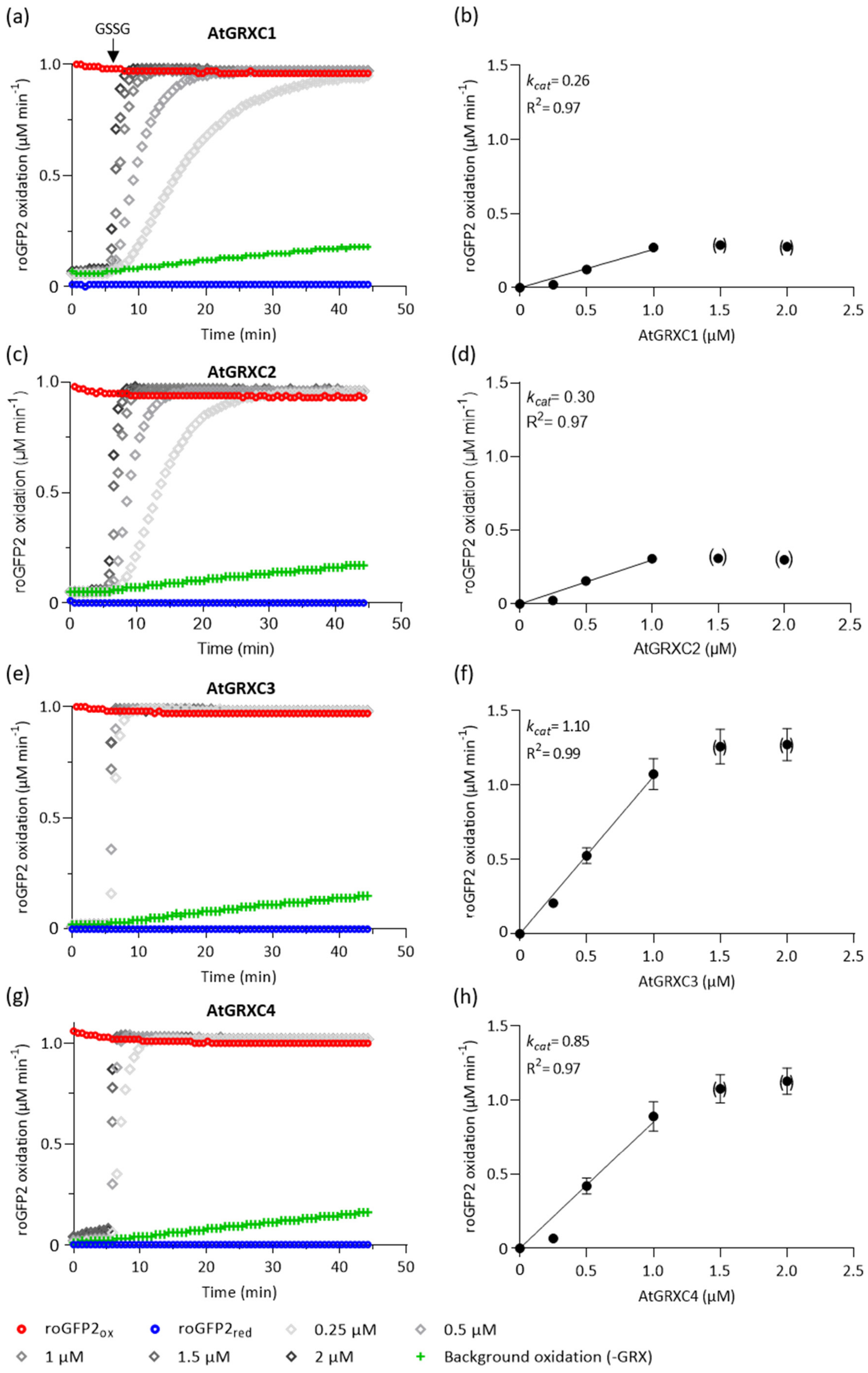

(a)
GSSG
AtGRXC1
roGFP2 oxidation (µM min-1)
Time (min)
(b)
kcat= 0.26
R2= 0.97
roGFP2 oxidation (µM min-1)
AtGRXC1 (µM)
(c)
AtGRXC2
roGFP2 oxidation (µM min-1)
Time (min)
(d)
kcat= 0.30
R2= 0.97
roGFP2 oxidation (µM min-1)
AtGRXC2 (µM)
(e)
AtGRXC3
roGFP2 oxidation (µM min-1)
Time (min)
(f)
kcat= 1.10
R2= 0.99
roGFP2 oxidation (µM min-1)
AtGRXC3 (µM)
(g)
AtGRXC4
roGFP2 oxidation (µM min-1)
Time (min)
(h)
kcat= 0.85
R2= 0.97
roGFP2 oxidation (µM min-1)
AtGRXC4 (µM)
roGFP2ox
roGFP2red
0.25 µM
0.5 µM
1 µM
1.5 µM
2 µM
Background oxidation (-GRX)

**Figure S11** Activity of class I GRXs in the oxidation of roGFP2. (a,c, e, g) GSSG dependent oxidation of 1 µM reduced roGFP2. All GRXs were tested from 0.25 µM (light grey) to 2 µM (dark grey). The background oxidation recorded the absence of GRX is depicted in green. Full reduction or oxidation was determined by incubating roGFP2 with 10 mM $DTT_{red}$ or 10 mM $DTT_{ox}$, respectively. The arrows mark the time point at which the assays were started by addition of 40 µM GSSG. (b, d, f, h) Concentrations of the GRXs used versus activity. After determining the reaction velocity for roGFP2 oxidation or reduction, the respective plots were fitted with a linear regression to extract $k_{cat}$ values. The points in parenthesis were excluded from the analyses. Data shown are mean ± SD for a single experiment with three technical replicates from one protein purification and dilution.

**TABLE S4** Kinetic properties of cytosolic and luminal class I glutaredoxins.

| Protein | HED assay | | | roGFP2 reduction | roGFP2 oxidation |
|---|---|---|---|---|---|
| | $K_m$ (µM) | $k_{cat}$ ($s^{-1}$) | $k_{cat}$ / $K_m$ ($M^{-1}$ $s^{-1}$) | $k_{cat}$ ($min^{-1}$) | $k_{cat}$ ($min^{-1}$) |
| **GRXC1** | 170 ± 57 | 17 ± 8 | 1 x $10^5$ | 0.09 ± 0.03 | 0.25 ± 0.03 |
| **GRXC2** | 397 ± 294 | 32 ± 20 | 0.9 x $10^5$ | 0.1 ± 0.01 | 0.38 ± 0.071 |
| **GRXC3** | 809 ± 434 | 131 ± 70 | 1.6 x $10^5$ | 0.04 ± 0.001 | 0.90 ± 0.17 |
| **GRXC4** | 1127 ± 653 | 219 ± 149 | 1.8 x $10^5$ | 0.04 ± 0.008 | 0.71 ± 0.16 |

***Note***: Data shown for the HED assay as mean ± SD, of four biological replicates with three technical replicates from different protein purification and dilution. For the roGFP2 assay the Data shown as mean ± SD, of three biological replicates with three technical replicates.

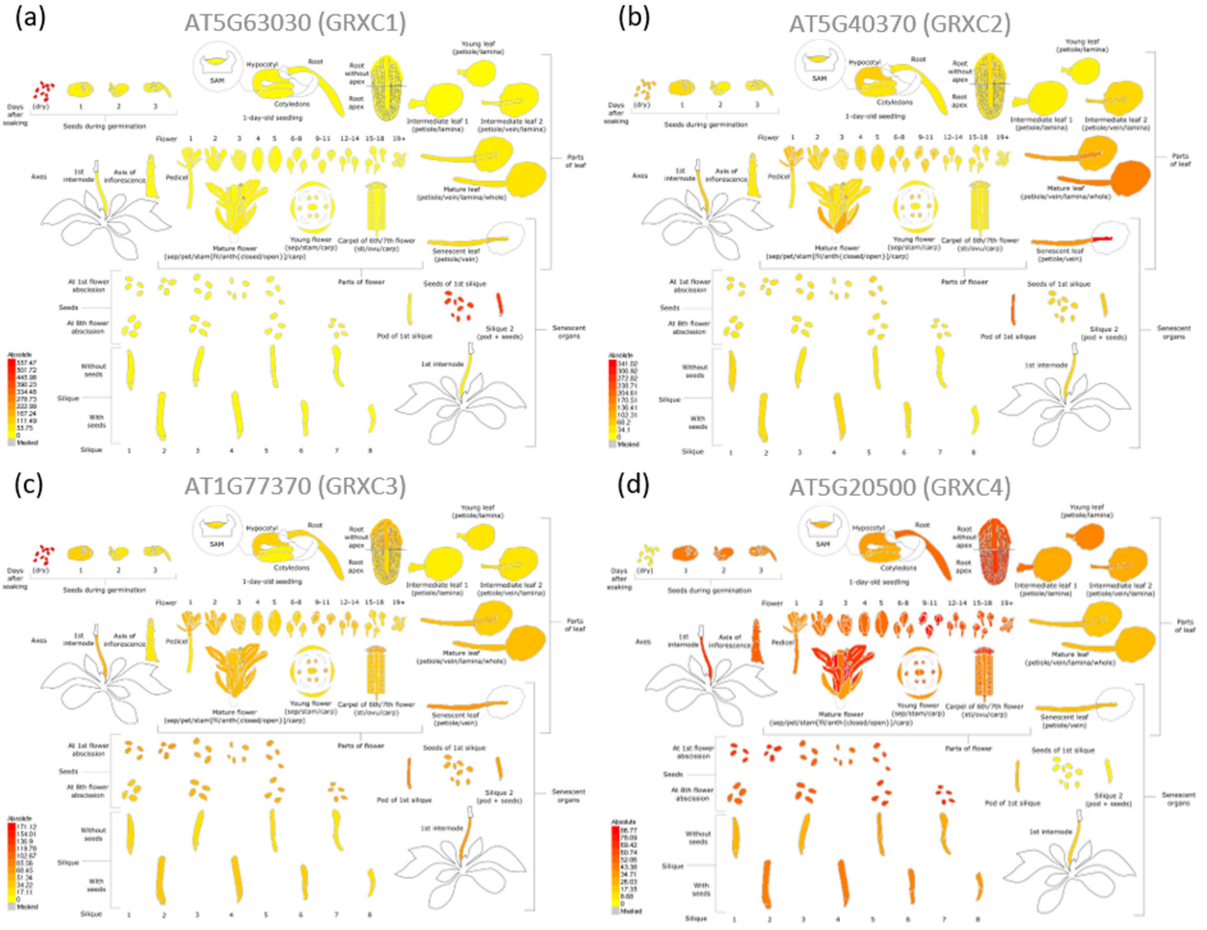


**Figure S12** Developmental map of expression for Arabidopsis GRXC1-4. Expression pattern of GRXC1 (a), GRXC2 (b), GRXC3 (c), and GRXC4 (d). The figures depict the absolute expression values based on data assembled in the Arabidopsis eFP Browser 2.0 (Winter et al., 2007).

WINTER D, VINEGAR B, NAHAL H, AMMAR R, WILSON GV, et al. 2007. PLOS ONE, 2(8): e718.
